# How to measure response diversity

**DOI:** 10.1101/2022.04.26.489626

**Authors:** Samuel R.P-J. Ross, Owen L. Petchey, Takehiro Sasaki, David W. Armitage

## Abstract

1. The insurance effect of biodiversity—that diversity stabilises aggregate ecosystem properties—is mechanistically underlain by inter- and intraspecific trait variation in organismal responses to the environment. This variation, termed *response diversity*, is therefore a potentially critical determinant of ecological stability. However, response diversity has yet to be widely quantified, possibly due to difficulties in its measurement. Even when it has been measured, approaches have varied.
2. Here, we review methods for measuring response diversity and from them distil a methodological framework for quantifying response diversity from experimental and/or observational data, which can be practically applied in lab and field settings across a range of taxa.
3. Previous empirical studies on response diversity most commonly invoke response traits as proxies aimed at capturing species’ ecological responses to the environment. Our approach, which is based on environment-dependent ecological responses to any biotic or abiotic environmental variable, is conceptually simple and robust to any form of environmental response, including nonlinear responses. Given its derivation from empirical data on species’ ecological responses, this approach should more directly reflect response diversity than the trait-based approach dominant in the literature.
4. By capturing even subtle inter- or intraspecific variation in environmental responses, and environment-dependencies in response diversity, we hope this framework will motivate tests of the diversity-stability relationship from a new perspective, and provide an approach for mapping, monitoring, and conserving this critical dimension of biodiversity.

## 1. Introduction

Ecological stability has been a core focus of ecology since the 1950s when interest in the relationship between diversity and stability arose (McCann 2000). Ecological stability is a multidimensional concept encompassing a variety of metrics for measuring resistance to and recovery from disturbance, and variability in time and space (Pimm 1984; Donohue *et al*. 2013). Early observations suggested that more diverse communities are more stable in their ability to resist invasions or in the magnitude of their fluctuations (Odum 1953; MacArthur 1955; Elton 1958). Using randomly structured model communities, May (1973) then demonstrated that diversity *per se* did not increase stability (that is, did not dampen population fluctuations), but rather decreased it (see also Pimm & Lawton 1978; Yodzis 1981). Subsequent theoretical developments (*e*.*g*., McCann *et al*. 1998; Kondoh 2003; Ives & Carpenter 2007) reached the consensus that species richness tends to stabilise aggregate ecosystem properties when complexity (in terms of, for example, variable environments, interaction strengths, or behaviours) is higher (Ives *et al*. 1999; but see Jacquet *et al*. 2016; Pennekamp *et al*. 2018). This is because declines in one species’ abundance or *performance* will be offset by neutral or positive responses of other species or phenotypes (reviewed by Loreau *et al*. 2021). We define performance as any measurable feature an organism or population that is an indicator of individual fitness. Functional traits that indicate performance could include both higher-level traits (such as per-capita growth rate) and lower-level traits (such as morphological traits) in the trait hierarchy (Agrawal et al. 2010). High-level traits are generally more likely to relate to environmental conditions than low-level traits (Violle et al. 2007). Together, these ideas encapsulate the insurance effect of biodiversity, which posits that biodiversity both enhances and stabilises ecosystem functioning (Yachi & Loreau 1999).

However, biodiversity is only the “passive recipient” through with underlying ecological mechanisms drive stability (McCann 2000; Shade 2017). Asynchronous or compensatory dynamics among species through time are now widely considered a key driver of the diversity-stability relationship (*e*.*g*., Craven *et al*. 2018; Sasaki *et al*. 2019; see also Loreau *et al*. 2021). Many specific biological mechanisms have been proposed as drivers of stability—largely through their effect on species asynchrony—including: variable interaction strengths (*e*.*g*., Yodzis 1981; McCann *et al*. 1998) or mixtures of interaction types (*e*.*g*., Mougi & Kondoh 2012; Hammill *et al*. 2015); food web nestedness (*e*.*g*., Thebault & Fontaine 2010); trophic flexibility and food web rewiring (*e*.*g*., Kondoh 2003); dominant, rare, or otherwise key species (*e*.*g*., Walker *et al*. 1999; Sasaki & Lauenroth 2011; Arnoldi *et al*. 2019; Ross *et al*. 2022a); and asynchronous patch dynamics in space (*e*.*g*., Loreau *et al*. 2003; Wang & Loreau 2016). Recently, the concept of *response diversity* has also emerged as a potentially key, yet under-explored driver of stability (Pennekamp *et al*. 2018; Kahiluoto *et al*. 2019; Sasaki *et al*. 2019).

Response diversity characterises the range of responses to the environment displayed among members (*e*.*g*., individuals, species) of a focal group such as a population, guild, or community and is *a priori* predicted to enhance community (aggregate) stability (Elmqvist *et al*. 2003; Nyström 2006; Mori *et al*. 2013). For example, Ives *et al*. (1999) posited that differences in how species respond to environmental fluctuations (that is, response diversity) may explain species diversity’s stabilising effect on total community biomass. In a later study, Ives & Carpenter (2007) found that, in a model of randomly structured competitive communities, response diversity’s stabilising effect on communities negated the destabilising effects of interspecific interactions. The authors concluded that species-environment relationships were more important for stability than were interspecific interactions (Ives & Carpenter 2007; see also Houlahan *et al*. 2007). Though such studies typically focus on species responses to fluctuating environments (*e*.*g*., Ives et al. 1999; Thibaut & Connolly 2013), species’ environmental responses are most easily characterised along environmental gradients (see Section 3). The importance of this distinction may be a matter of scale. The rate at which organisms experience environmental change impacts their physiological and ecological capacity to respond to such change (Wolkovich et al. 2014; Pinek et al. 2020), including the likelihood of local adaptation along a gradient and under temporally variable environmental conditions (*e*.*g*., Villa Martín et al. 2019). The perceived rate of environmental change is itself a consequence of life history: what one short-lived organism perceives as a gradual (or abrupt) environmental change may simultaneously be perceived as a temporary environmental flux by another, longer-lived organism (Wolkovich et al. 2014; Jackson et al. 2021). Measuring species-environment relationships over environmental gradients is therefore practically useful for informing about species and community responses to temporal environmental fluctuations.

Interspecific variation in species-environment relationships is critical for the maintenance of diversity in variable environments (Chesson 2000) and species-rich systems are thus anticipated to exhibit a wider diversity of species-environment interactions compared to species-poor systems (Vogel *et al*. 2019). This response diversity may, in turn, support a variety of stabilising mechanisms acting on the aggregate properties of communities and ecosystems. For example, response diversity may be one candidate ecological mechanism underpinning many of the statistical explanations of the diversity-stability relationship, such as the portfolio effect (Tilman *et al*. 1998; Loreau 2010; Loreau *et al*. 2021), particularly through its effect on species asynchrony. Response diversity can promote asynchrony in species’ biomass or abundance through time (Sasaki *et al*. 2019) and species-specific tolerance to environmental conditions may be partly responsible for temporal and functional complementarity among species (Petchey 2003; Loreau & de Mazancourt 2013). Indeed, asynchrony may frequently be an outcome of response diversity (*e*.*g*., Sasaki *et al*. 2019), suggesting that the often-observed stabilising effect of species asynchrony on aggregate community or ecosystem properties (*e*.*g*., Morin *et al*. 2014; Hector *et al*. 2015), may result from underlying differences in species-environment relationships. It should therefore benefit those interested in the diversity-stability relationship to measure both species asynchrony and response diversity—given that response diversity should predict asynchrony, and in turn stability—as well as any additional factors expected to drive community stability that may be of interest, such as species richness *per se*, spatial synchrony, or population stability.

## 2. Existing response diversity studies

Few empirical studies have explicitly measured response diversity. Recognising that many studies may measure response diversity without labelling it such, a systematic review of the literature returned 46 ecology papers that empirically measured response diversity (Supporting Figure S1). These empirical studies used various methods to measure response diversity. The majority (*n* = 28 studies, ∼60%) used low-level functional traits (see Box 1 for discussion), where the diversity of response traits—that is, those traits that predict some aspect of how a species responds to the environment (Suding *et al*. 2008)—represents response diversity, measured using the functional dispersion [FDis] index (*e*.*g*., Spasojevic *et al*. 2016; Hordley *et al*. 2021; Schnabel *et al*. 2021). Some studies measuring the dispersion of response traits explicitly do so after defining functional effect groups (Laliberté *et al*. 2010; Thornhill *et al*. 2018; Sasaki *et al*. 2019; Morel *et al*. 2020) following Elmqvist *et al*. (2003), though subsequent definitions relax this requirement (*e*.*g*., Mori *et al*. 2013). Most empirical studies measuring response trait diversity do so for trees (*e*.*g*., Craven *et al*. 2016; Altomare *et al*. 2021; Schnabel *et al*. 2021) and other groups of plants (*e*.*g*., Mandle & Ticktin 2015; Döbert *et al*. 2017; Morel *et al*. 2020), or for freshwater or terrestrial invertebrates (*e*.*g*., Mumme *et al*. 2015; Thornhill *et al*. 2018).

Another approach to measuring response diversity empirically is to define a species-specific interaction term in multi-species models of, for example, abundance as a function of the environment (*n* = 7 studies; Figure S1). This binary approach asks whether there is response diversity among species or not. If the species-specific interaction term passes some significance or model selection threshold, then species are claimed to differ in their abundance ∼ environment slopes; that is, they exhibit response diversity (Winfree & Kremen 2009). This approach to measuring response diversity has emerged primarily from the pollination literature (Winfree & Kremen 2009; Bartomeus *et al*. 2013; Cariveau *et al*. 2013; Stavert *et al*. 2017), but has been used elsewhere (Fauchald *et al*. 2011; Malyshev *et al*. 2016). Another related but less common approach to measuring response diversity is to measure some aspect of a population’s performance, such as its intrinsic rate of increase (*e*.*g*., McCann 2016) or biomass change in response to the environment, and take the range of performance-environment model slopes as a direct estimate of response diversity. Leary and Petchey (2009) did this with four ciliate species by modelling their intrinsic rates of increase against temperature when in isolation, first determining species-specific environmental responses. They then assembled experimental communities with different pairwise combinations of these species, producing different levels of response diversity. A conceptually similar approach from Thibaut *et al*. (2012) used observational time series of reef fish abundance to measure response diversity based on the average correlation coefficient of random effects (intercept terms) on the intrinsic rates of increase between functional groups. Unlike Leary and Petchey’s (2009) approach for considering pairwise performance-environment correlations, this approach directly quantifies covariation among responses to environmental fluctuations using pairwise correlations in the deviation from the average intrinsic rates (*i*.*e*., the random effect term; Thibaut *et al*. 2012). These various methods for measuring response diversity (Figures 1 and S1) have emerged independently, resulting in a disconnect among empirical response diversity studies, and a general methodological framework for measuring response diversity is lacking.

**Figure 1.**
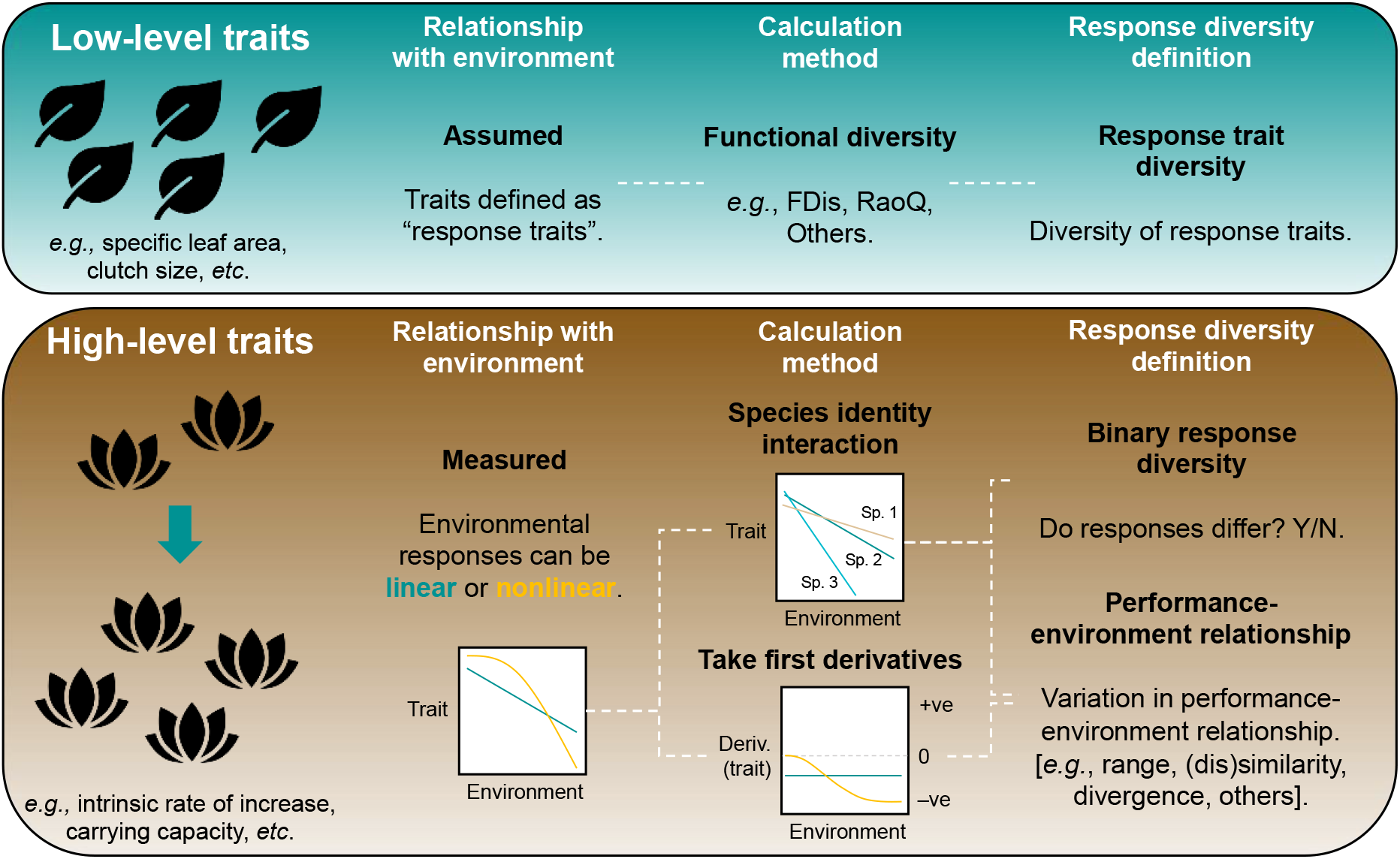
How to measure response diversity. Response diversity can be measured from low-level functional traits—the approach dominant in the literature (Figure S1)—or from higher-level traits which are themselves sometimes underlain by low-level traits. One limitation of measuring response diversity based on low-level traits is that such approaches often assume relationships between traits and the environment, rather than directly measuring trait-environment relationships (Box 1). Some previous studies have directly measured trait-environment relationships and tested whether different species respond differently to the environment (“species identity interaction”; Section 2). Such measurements can be linear or nonlinear. We propose taking the first derivative of such trait-environment relationships to better account for nonlinear environmental responses (Section 3.2). One can then determine whether all species respond similarly to the environment (“binary response diversity”; Section 2) or can go further and measure aspects of interspecific variation in trait-environment relationships (*i*.*e*., the “performance-environment relationship”; Section 3) using methods described here or elsewhere.

In response to this, in the remainder of this article we present a framework for measuring response diversity empirically using performance-environment relationships (Section 3), including application to simulated data for linear (Section 3.1) and nonlinear species responses where the form of the performance-environment relationship depends on environmental context (Section 3.2), as well as an application to an empirical dataset of multispecies aquatic microcosms (Section 3.3). Finally, we discuss future directions for empirical studies of response diversity and the challenges of applying this approach in a range of contexts (Section 4).

## 3. Measuring response diversity empirically

Here, we propose a framework for empirically measuring response diversity using a conceptually simple and empirically tractable approach that can be applied in observational and experimental settings in both the laboratory and field. We extend the models describing performance-environment relationships presented above to include any response variable that represents a species’ ecological response along a continuous gradient of abiotic or biotic environmental condition. In contrast to most previous studies of response diversity which focus on ‘low-level’ functional traits such as nitrogen content or specific leaf area (Box 1), our approach considers response diversity a ‘high-level’ trait of *per capita* growth rates for instance, tying it more directly to species performance (see Violle *et al*. 2007; Agrawal *et al*. 2010). That said, low-level traits can still be used in this framework, and can in some cases be informative and important responses (*e*.*g*., stand-level Nitrogen content).

Our aim is not to define a single best metric of response diversity for use in all cases, but rather to illustrate how it can be measured. We demonstrate two complimentary metrics with desirable properties for our purposes. More suitable metrics for different use-cases likely exist, and we encourage researchers to explore various methods while warning against the proliferation of untested response diversity metrics—a new metric should necessarily be an improvement on existing metrics in its ability to fulfil its purpose (see Figure 5 below). In response diversity’s case, this purpose is to predict ecological stability, as grounded in the basic principles of the insurance hypothesis (Yachi & Loreau 1999), while noting that response diversity may not always be the only driver of stability or species asynchrony.

We propose extending the performance-environment models used elsewhere (Leary & Petchey 2009) into a generalisable framework for measuring response diversity. Despite rarely being used to date, this method is easily tractable in experimental and observational studies, and provides a more direct estimate of how species respond to the environment than, for example, the often-used response trait approach (Box 1). Given our definition of *performance*, the response variable in such models need not be intrinsic rate of increase, but could in fact be any relevant functional variable, such as biomass, abundance, or an ecosystem function to which a species contributes. Similarly, the *environment* to which species are responding does not have to be an abiotic feature such as temperature; it can be any continuous biotic or abiotic condition. For example, one could model the responses along a land-use intensification gradient (Winfree & Kremen 2009; Moore & Olden 2017; Stavert *et al*. 2017), or in response to the abundance of a predator, competitor, or the availability of a prey species (*e*.*g*., Fauchald *et al*. 2011). The latter approach, using biotic conditions as the environment to which a species responds, is conceptually similar to modelling the functional response (Holling 1966), which in turn may scale with abiotic conditions such as temperature (Daugaard *et al*. 2019). Likewise, modelling biomass change in response to a competitor is akin to concepts in coexistence theory such as negative density-dependence (*e*.*g*., Armitage & Jones 2019) or invasion growth rate (Grainger *et al*. 2019).

### 3.1. Measuring variation among performance curves

From such models, we suggest measuring response diversity as the variation in the slope (first derivative) of a performance-environment function evaluated over the environmental gradient in question. Specifically, we suggest using Generalised Additive Models (GAMs) to fit performance-environment relationships individually for each species using smoothing parameters as appropriate (Figures 2a-2c), then taking the first derivatives of these models to estimate model slopes along the environmental axis (Figures 2d-2f). Note that it may not always be necessary to calculate the derivatives of a performance variable that is already a rate measure; in this case, the fitted GAM function would suffice. GAMs are free from many of the assumptions of linear regressions, such as normality and homoscedasticity of residuals, and can be used to evaluate both linear and nonlinear responses, or a mixture of the two (Hastie & Tibshirani 1987; see Ross *et al*. 2022b). Response diversity can then be measured as the variation in the distribution of these derivatives to capture the variation in species-specific environmental responses (Figures 2g-2i). In practice, this variation could be described using any suitable estimator (Figure 1), but here we use two complementary metrics.

**Figure 2.**
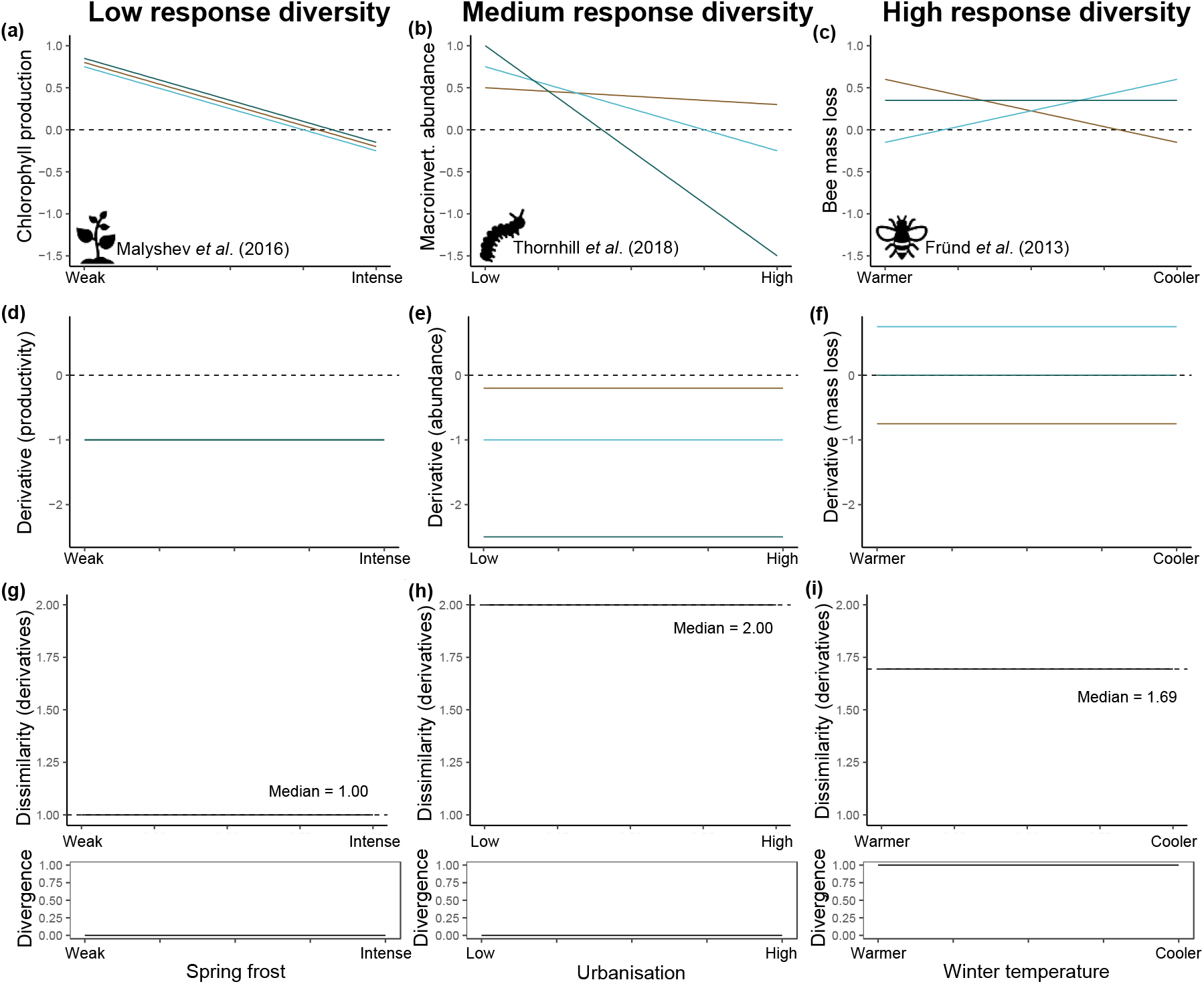
Measuring response diversity for linear species-environment relationships. We reimagined the original analyses of three studies, using standardised, simulated data to demonstrate how response diversity could be measured practically under our framework. **(a)** We proposed a hypothetical case (based on Malyshev *et al*. 2016), where *Chlorophyll* production (which is often an indicator of biomass, particularly for aquatic plants) declines linearly across several species [coloured lines] of plants with spring frost intensity, with no difference among species in the slope of this decline. **(d)** We calculated the first derivatives of these responses, which in this case was identical for all species. **(g)** Then, we measured response diversity using the proposed similarity-based diversity metric (see main text), and found it to be one (its lower bound), since all species respond identically to spring frost. Our divergence metric was zero in this case, as the first derivatives do not span zero. **(b)** Next consider a case (based on Thornhill *et al*. 2018), where the abundance of several species of macroinvertebrates declines linearly, but at different magnitudes, along an urbanisation gradient. **(e)** Here, the first derivatives differed (owing to differing model slopes), resulting in higher dissimilarity **(h)**. Again, the derivatives did not span zero, so divergence was zero. **(c)** Finally, we might be interested in the decline in mass of several bee species as a function of winter temperature (see Fründ *et al*. 2013). In this example, species responses to winter temperature diverged (one species increased, one decreased, one was unaffected), but all such responses were linear, and the magnitude of responses was not as extreme as in **(b). (f)** Thus, the first derivatives were less spread out than in **(e)**, but spanned zero. **(i)** Accordingly, dissimilarity was lower than for **(h)**, but in this case, divergence was at its maximum value, 1, across the entire range of winter temperature values (the derivatives spanned zero at any temperature). Dashed lines indicate zero in panels **a–f**, and the median value of the similarity-based diversity metric (dissimilarity) along the X-axis in panels **g–I** (here, equivalent to any value along the X-axis since response diversity is constant).

First, we use a similarity-scaled metric of diversity introduced by Leinster & Cobbold (2012). Briefly, similarity-based diversity (Eq. 1 in Leinster & Cobbold 2012) is calculated based on pairwise Euclidean distances (that is, *dissimilarity*) in performance-environment relationships between all pairs of species in the community. The metric is well suited to measuring response diversity, since it combines similarities and relative abundances, with an exponent (*q*) tuning the weighting placed on relative abundance (Leinster & Cobbold 2012). Here, we fix *q* at zero, to focus solely on species richness (*S*), though this (dis)similarity metric can use information on species relative abundances within an *S* x *S* similarity matrix. We chose this estimator since it has several desirable properties for describing response diversity: it cannot decrease with species additions; does not increase with additions of redundant species responses; and captures even subtle differences in the distribution of different responses within the overall range. Dissimilarity should therefore be lowest (1) when all species respond identically (Figure 2g), and highest (*S*) when species responses are as dissimilar as possible (the estimator converges to species richness as performance-environment slopes diverge to infinity and –infinity).

Second, we develop a simple measure that accounts for whether performance-environment responses *diverge* in direction, since this should be a key indicator of the stabilising potential of response diversity under a given environmental condition. The insurance effect of biodiversity assumes that declines of one species can be offset by neutral or positive responses of others (Yachi & Loreau 1999). As such, we also isolate and measure this property of response diversity using the formula:

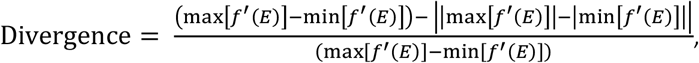

where *f*′(*E*) represents the first derivative of a performance-environment relationship, *f*(*E*), at a particular environmental state (*E*). If the range of derivatives does not span zero (*e*.*g*., Figures 2d and 2e), then all species respond in the same direction to the environment (albeit with varying model slopes). In such cases, there is no response diversity along this divergence axis of species responses (the value is zero). Without interspecific interactions, even large differences in the magnitude of response (dissimilarity) cannot be stabilising (Ives *et al*. 1999; Mougi & Kondoh 2012). In contrast, when derivatives are perfectly symmetric around zero, the value is one. When the range of first derivatives spans zero for a given environmental state (*e*.*g*., Figure 2f), species responses are diverging (at least one species is decreasing while another is increasing), which we expect *a priori* to be stabilising. This divergence metric is largest (1) when min[*f*′(*E*)] = −max[*f*′(*E*)]. Observation of stable community biomass or other aggregate ecosystem properties when the range of performance-environment derivatives does not span zero would suggest that divergence in species responses is not the mechanism driving stability in that case.

Together, the similarity-based diversity metric and divergence metric make up two complimentary components of response diversity; the similarity-based metric captures the overall magnitude of response diversity among species’ responses (Leinster & Cobbold 2012), while the divergence metric aims to quantify the extent to which species declines are offset by gains (Yachi & Loreau 1999). Though often independent, these two metrics may respond similarly to changes in species’ performance-environment relationships. For example, when mean responses (derivatives) are close to zero, an increase in dissimilarity will necessarily cause an increase in divergence. The specific metrics we use here may not be ideal for all use cases. Further work should consequently consider the need to develop suitable metrics of response diversity when those proposed fall short. There may also be opportunity to unite the two dimensions of response diversity discussed here into a single metric. For example, when (dis)similarity and divergence are highly correlated—such as when performance-environment relationships are nonlinear and divergence is high (Figure 3)—the simultaneous use of both metrics does not provide much additional information on response diversity over one metric in isolation (see Donohue *et al*. 2013), so using a single metric may suffice. That said, we demonstrate here the utility of these metrics for quantifying response diversity in a range of contexts (Figures 2 and 3), including applications to real data (Figures 4 and 5).

**Figure 3.**
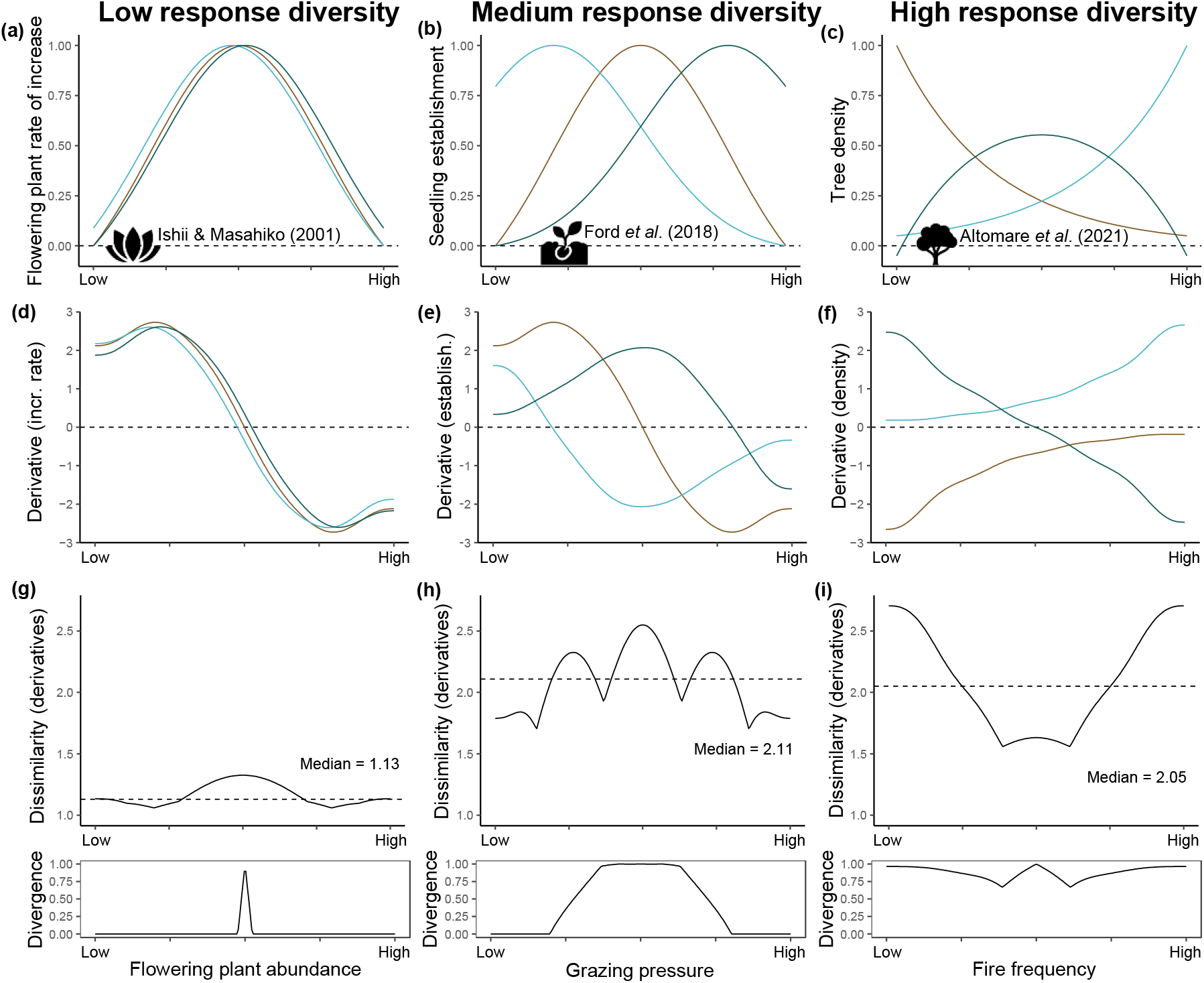
Measuring response diversity for nonlinear species-environment relationships. We reimagined the original analyses of three studies, using standardised, simulated data to demonstrate how response diversity could be measured practically under our framework. **(a)** We proposed a hypothetical case (based on Ishii & Masahiko 2001), where flowering plant intrinsic rate of increase changed as a function of total flowering plant abundance. Here, responses were nonlinear but similar in form, as competition was limiting at high abundance while pollinator visitations were perhaps limiting where total flowering plant abundance was low (Ishii & Masahiko 2001; Reilly *et al*. 2020). **(d)** In this case, the first derivatives of the intrinsic rates of increase varied as a function of flowering plant abundance, but did not differ much among species. **(g)** This resulted in a dissimilarity value close to its minimum (1) but that varied a little with total flowering plant abundance. **(b)** Next consider a case (based on Ford *et al*. 2018), where seedling establishment of several plant species responded nonlinearly to grazing pressure. This might be the case if, for example, one species is a shade-tolerant understorey specialist [light blue] that does well when grazing pressure is low but not when high, while another species is a shade-intolerant disturbance specialist [dark blue] (Ford *et al*. 2018). **(e)** These divergent nonlinear responses produced a range of first derivatives that changed as a function of grazing pressure, in turn producing **(g)** grazing pressure-dependent response diversity which was on, average, high in terms of dissimilarity, and was zero-spanning at intermediate grazing pressure. **(c)** Finally, consider a case (based on Altomare *et al*. 2021), where the density of several tree species responded nonlinearly to fire frequency with different functional forms. This might occur if some species were highly sensitive to burning [brown line] while others were disturbance specialists that require nutrient inputs from fire or reduced competition from dominant fire-sensitive species (Altomare *et al*. 2021). **(f)** Here, the first derivatives of tree density differed considerably and varied as a function of fire frequency, producing **(i)** generally high response diversity—in both dissimilarity and divergence—that also varied with fire frequency. Here, response diversity was largest at low and high fire frequency (when specialists responded strongly) and was lowest at intermediate fire frequency. Dashed lines indicate zero in panels **a–f**, and the median value of the similarity-based diversity metric (dissimilarity) along the X-axis in panels **g–i**. Note that as average divergence increases from **g** to **i**, the Pearson correlation coefficient (*r*) between dissimilarity and divergence increases: **(g)** Pearson’s *r* = 0.36, **(h)** *r* = 0.70, **(i)** *r* = 0.75.

**Figure 4.**
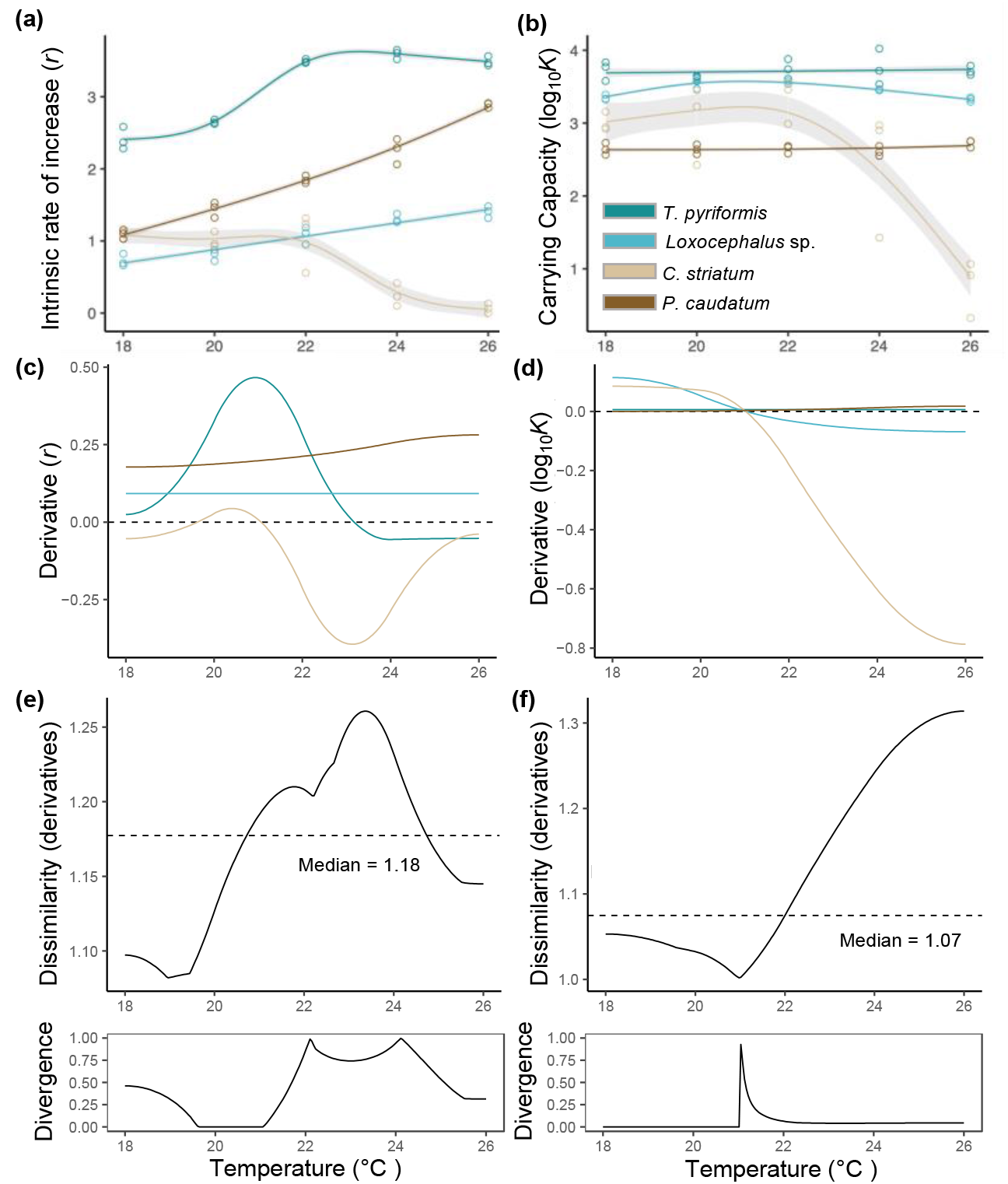
Re-analysis of data from Leary and Petchey (2009) using the framework proposed herein. Using empirical data from Leary and Petchey (2009), we measured performance-environment relationships for four aquatic ciliate species [coloured lines] as **(a)** the intrinsic rate of increase [*r*] and **(b)** the carrying capacity [log_10_*K*] along a temperature gradient (18-26°C). Data points are from a single population (see Figures 1a and 2a in Leary & Petchey 2009). **(c**,**d)** We calculated the first derivatives of the performance-temperature relationships, which changed as a function of temperature due to nonlinear responses. **(e**,**f)** We then measured the similarity-based diversity and divergence of the derivatives of all species in the community for both **(e)** *r* and **(f)** log_10_*K*. **(e)** For *r* we found that both indices peaked around 22-24°C. **(f)** For log_10_*K*, dissimilarity peaked at the upper temperature limit, as the response of one species (*C. striatum*) diverged from the others **(b**,**d)**, while divergence was mostly zero across the temperature range. Dashed lines indicate zero in panels **c** and **d**, and the median value of the similarity-based diversity metric (dissimilarity) along the X-axis in panels **e** and **f**.

**Figure 5.**
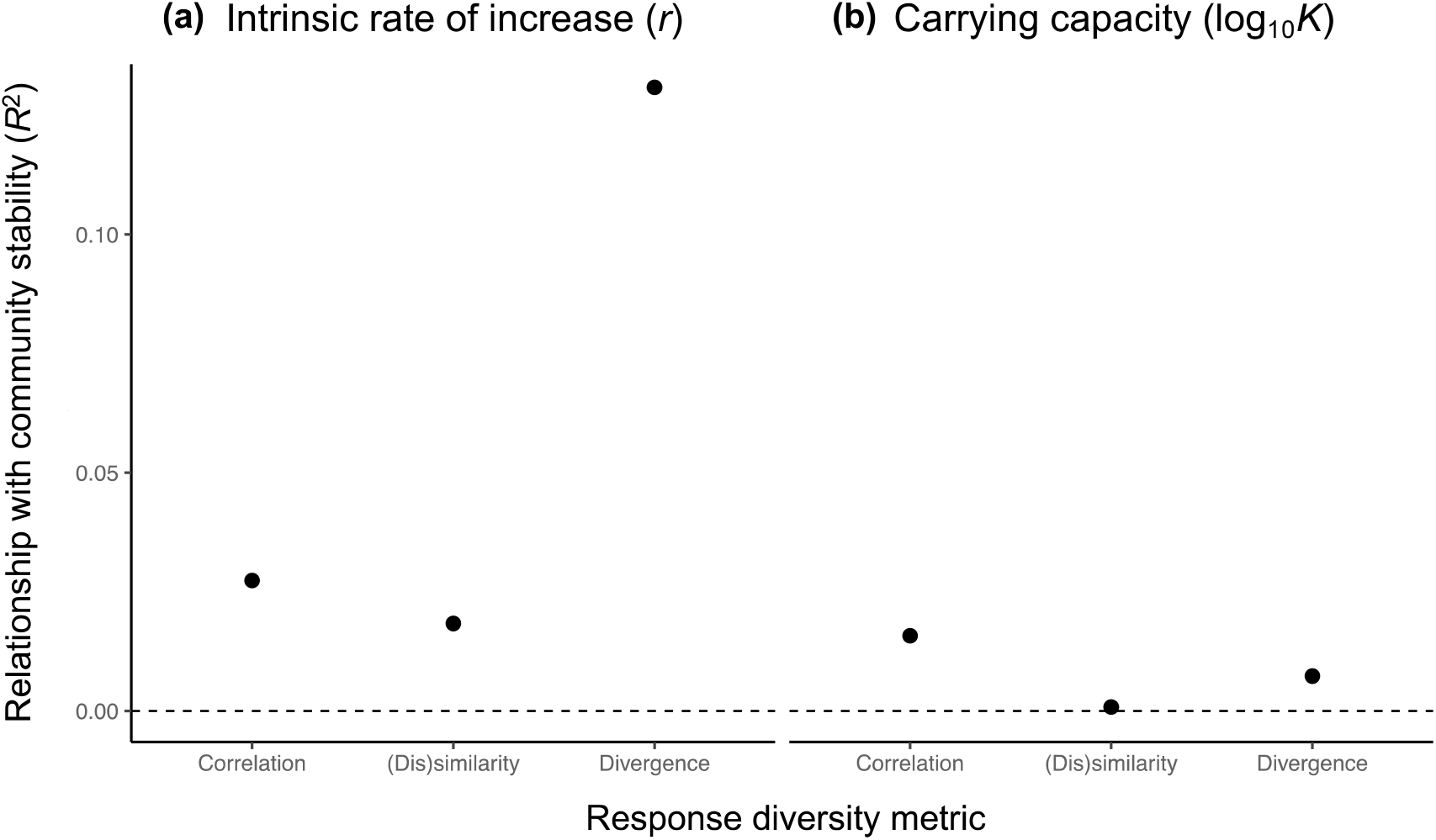
Comparison of response diversity-stability relationships for different metrics for data from Leary and Petchey (2009). After measuring response diversity for Leary and Petchey’s (2009) ciliate dataset using the framework proposed herein, we compared the ability of response diversity to predict community stability (the coefficient of variation of total biomass for each pairwise combination of ciliate species; see Leary & Petchey 2009; Ross *et al*. 2022b). Relationship with community stability is approximated by the predictive power (*i*.*e*., goodness-of-fit) using *R*^2^ of the response diversity-stability relationship, where higher *R*^2^ values represent higher predictive power. We tested three measures of response diversity: 1) a pairwise correlation-based measure similar to Leary and Petchey (2009) (*Correlation* on x-axis); 2) the Leinster & Cobbold (2012) similarity-based measure described therein (*(Dis)similarity*; Ross *et al*. 2022b); and 3) the divergence metric measuring the proportion of zero-spanning species’ derivatives (*Divergence*). Panel **(a)** represents response diversity and community stability measured from the Intrinsic rate of increase, while **(b)** is measured from the log_10_ carrying capacity (Leary & Petchey 2009; Ross *et al*. 2022b). See Ross *et al*. 2022b for figures showing individual response diversity-stability relationships and a complementary *R*^2^ figure relating response diversity to stability calculated using covariance rather than coefficient of variation. Note that while *R*^2^ was highest for the Divergence metric for the intrinsic rate of increase **(a)**, none of the *R*^2^ values here were particularly high, suggesting that even our best performing index did not predict community stability especially well.

### 3.2 Nonlinear responses to environmental variation

Ecological responses to the environment are often nonlinear, necessitating a wide range of statistical tools for considering such responses. When species’ responses are nonlinear (*e*.*g*., Leary & Petchey 2009), quantifying response diversity is not as simple as measuring the range of model slopes. However, the statistical framework we propose for measuring response diversity (that is, measuring variation in performance-environment derivatives) is robust to nonlinearities such as species’ responses to competition, predation, or resource availability (Figure 3) or when tipping points are detected in empirical data (Berdugo *et al*. 2020). Figure 3 illustrates three such cases: one where the intrinsic rate of increase of flowering plant species responds nonlinearly but with low response diversity to total flowering plant abundance (Figure 3a); one where seedling establishment responds to grazing pressure with higher dissimilarity among species responses (Figure 3b); and one where trees respond nonlinearly to fire frequency, and diverge in their responses (Figure 3c). Our framework uses Generalised Additive Models as they are robust even to cases where the form of the performance-environment relationship differs among species (Hastie & Tibshirani 1987; Figures 3f and 3i). Where responses are nonlinear, the derivatives of performance-environment responses vary as a function of the environmental variable (along the X-axis in Figures 3d-3f). In turn, this produces estimates of dissimilarity and divergence that also vary with the environment (Figure 3g-3i). Using these hypothetical examples, we see that the performance variables of interest should be stable only over a narrow range of environmental conditions in the case of flowering plant abundance (Figure 3g), whereas seedling establishment and tree density are predicted to be stable over a wider set of environmental conditions (Figures 3h and 3i). In cases where there is no need to treat response diversity as environment-dependent, one could estimate a single response diversity value across the environmental axis using any summary statistic (see dissimilarity in Figures 3g-3i), though it may be appropriate to weight such a summary statistic by the distribution of the environmental variable. In linear cases, response diversity does not vary with environmental context, so summary statistics are not needed (Section 3.1).

By modelling individual species responses to environmental conditions with GAMs, this approach is conceptually similar to the binary species-specific interaction term described above (Winfree & Kremen 2009; Cariveau *et al*. 2013; Figure 1). However, the framework we describe provides a quantitative estimate of response diversity, identifying precisely the relevant differences in the form of species environmental responses (dissimilarity and divergence) that should drive ecological stability as described under the insurance hypothesis (Yachi & Loreau 1999), and allowing consideration of response diversity as a function of the environment. This method is flexible enough to be used on traits, rates (first derivatives), or sensitivities of rates (second derivatives). We demonstrate cases taking first derivatives of traits before calculating response diversity (*e*.*g*., Figure 2b), thus producing *rates* from which we calculate response diversity (Figure 2e). However, it is also possible to estimate derivatives of rates (*e*.*g*., Figures 3d and 4), producing a measure of sensitivity of those rates to environmental change, which can be ecologically meaningful when measured over environmental gradients (Shyu & Caswell 2014). However, study designs often estimate intrinsic rates of increase, for instance, under only two environmental contexts, such as in disturbed versus in-tact habitats. In such cases, it may be more meaningful to estimate response diversity based directly on the intrinsic rates of increase (rather than transforming them to second derivatives of abundance).

Individual use cases will differ, so we suggest researchers carefully consider whether response diversity should be measured based on traits, rates, or sensitivities, depending on available data and study goals (Figure 1). Practical users of response diversity can make use of the mgcv and gratia packages in *R* to fit GAMs and calculate derivatives, respectively (Wood *et al*. 2016; Simpson 2021), and we provide *R* code to calculate dissimilarity and divergence for the simulated case studies from Figures 2 and 3 and the reanalysis presented below (available through Github: Ross *et al*. 2022b).

Using the framework for measuring response diversity we present here, one can measure response diversity empirically in the laboratory or field. A conceptually simple and empirically tractable metric of response diversity can stimulate research on a range of previously unanswerable questions that can be experimentally tested in controlled systems. For example, once the range of response trajectories among species—or individuals/populations for cases of intraspecific response diversity—is established, communities encompassing differing amounts of response diversity can be manually assembled (Leary & Petchey 2009; Baskett *et al*. 2014). In turn, this allows testing a variety of questions with response diversity (either dissimilarity or divergence) as the predictor variable, such as whether experimentally assembled communities with higher response diversity are more stable in terms of, for example, the temporal variability of total community biomass. Such stability analyses will require repeated temporal sampling of aggregate community biomass or ecosystem functioning. This then necessitates a decision about when to measure response diversity if analysing its relationship with temporal stability (see discussion in Box 1).

### 3.3 Revisiting the response diversity-stability relationship in experimental aquatic communities

Next, as an empirical proof-of-concept for our framework, we demonstrate how it can be used to measure response diversity among aquatic microorganisms first presented by Leary and Petchey (2009). The original study involved four ciliate species (*Colpidium striatum, Loxocephallus* sp., *Paramecium caudatum*, and *Tetrahymena pyriformis*). The first empirical step was to measure the responses of each species alone to temperature (five levels: 18, 20, 22, 24, and 26°C; Figures 4a and 4b). Responses of two types of performance were estimated from population time series: intrinsic rate of increase (*r*) and carrying capacity (log_10_ *K*). These two properties (*r* and log_10_*K*) are uncorrelated here (Spearman’s *rho* = 0.129) and provide complementary information on species’ performance, each with relevance at different temporal scales. For example, *r* is likely of more relevance if the environmental change of interest occurs rapidly or is short-lived, since *r* provides information on a population’s ability to rapidly bounce back after disturbance. Conversely, log_10_*K* may be most relevant over longer temporal scales of environmental change, since populations are unlikely to reach carrying capacity immediately following disturbance. The second empirical step was to record the biomass dynamics of communities that contained different combinations of the four species, and which were exposed to different ranges of temporal variation in temperature. The original analyses involved calculating the response diversity present in each of the communities, and comparing this to the stability of the biomass dynamics of the communities (Leary & Petchey 2009).

Here, we took a different approach to measure response diversity than in the original study. We modelled each species’ performance (*r* or log_10_*K*) individually using Generalized Additive Models (GAMs) and calculated the first derivates of each performance-temperature relationship (Figures 4c and 4d). We then estimated the two dimensions of response diversity described above: the (dis)similarity of first derivatives (Leinster & Cobbold 2012) and their divergence (the extent to which derivatives span zero for a given temperature). In doing this as a function of temperature, we produced temperature-dependent estimates of dissimilarity and divergence, which were highest for *r* at around 23.5°C when performance-environment responses were divergent and zero-spanning (Figure 4e), and highest for log_10_*K* at the maximum temperature as the carrying capacity of *C. striatum* declines while the other species remain fairly constant (suggesting *C. striatum* has a disproportionate contribution to dissimilarity here). Note, however, that the divergence of the derivatives is low at higher temperatures for log_10_*K*, so we might predict any community-level properties driven by carrying capacity to be unstable at high temperatures (Figure 4f). Such temperature-dependent response diversity values provide an added layer of information—and hence predictive capacity— compared to the original analyses based on correlated performance-environment responses (Leary & Petchey 2009).

Just as metrics measuring functional diversity with effect traits should be related to ecosystem functioning (Cadotte *et al*. 2011), response diversity measures should explain and/or predict community stability. Next, we assess this explanatory power, again using Leary and Petchey’s (2009) aquatic ciliate experiment. To do so, we compared the relationship between response diversity and community stability of pairs of ciliate species. Stability is defined here as the coefficient of variation (ratio of the standard deviation to the mean) of aggregate ciliate biomass for each species pair (Donohue *et al*. 2013).

We calculated six measures of response diversity—three metrics, each applied to the *r* and log_10_*K* response curves. We calculated the two metrics of response diversity discussed above: the Leinster and Cobbold (2012) similarity-based measure, and the divergence measure aimed at capturing whether species responses differ in their direction. We also calculated response diversity following the original study; we measured the correlation between the *r* or log10*K* responses estimated for the species in each community. To assess the performance of different measures of response diversity, we then modelled the relationship between each response diversity metric and community stability using a simple linear regression. Specifically, for each species pair (see Leary & Petchey 2009), we used Pearson correlations between each response diversity metric separately and the coefficient of variation of aggregate (two species) community biomass (using summed covariances as the stability measure produced similar results; Ross *et al*. 2022b). Where model slopes (± 95% confidence intervals [C.I.]) did not span zero, we interpreted this as a significant relationship (positive or negative) with stability (Figure 5). This approach, based on the regression models from Leary and Petchey (2009) provides only a first and illustrative test of the performance of each response diversity metric. More sophisticated models of the relationship between response diversity and stability could likely account for some dependencies among observations, or for nonlinear relationships between response diversity and stability, which need not be simple linear functions (Ives *et al*. 1999). To concretely demonstrate that a given response diversity metric performs better than any other thus likely requires *a priori* information on the expected form of the response diversity-stability relationship. Using simulations to test metric performance for known response diversity-stability relationships will likely be an important step in validating new metrics. Nevertheless, our approach based on simple linear regression provides a first proof-of-concept for linking response diversity measured from performance-environment relationships to ecological stability and assessing the relative performance of different response diversity metrics.

We found that for the intrinsic rate of increase, our proposed divergence metric outperformed other metrics of response diversity (including that from the original study) in terms of the predictive power (*R*^2^) between response diversity and aggregate community biomass stability (Figure 5a). This metric therefore represents an advance when aiming to predict community stability of *r*, though the Leinster and Cobbold (2012) method was not a significant improvement, and divergence did not improve the relationship with log_10_*K* stability (results of covariance-based stability were again similar; Ross *et al*. 2022b). Future efforts should thus use this example as a springboard from which to explore a wider range of response diversity methods, using data from various case studies, with the explicit goal of benchmarking any proposed new response diversity metric against existing approaches to determine which method best predicts community stability. In doing so, methodological development should not deviate from response diversity’s *a priori* theoretical aim to reflect the diversity of species responses to the environment and, in turn, to mechanistically drive ecological stability via asynchrony (Yachi & Loreau 1999).

## 4. Future directions and challenges

The framework for measuring response diversity introduced above is conceptually simple and applicable across contexts. Using this approach, response diversity can be easily measured in empirical settings (see Figure 4 for reanalysis of data from Leary & Petchey 2009), experimentally manipulated, and related to stability where suitable community time series are available (Figure 5). This offers advantages over the functional trait approach dominant in the literature which estimates species’ environmental responses less directly (Box 1 and Figure 1)—though future work should make direct comparisons between response diversity measured using performance-environment relationships and those estimated using low-level functional traits.

### 4.1. Interspecific interactions

The performance-environment relationships we mention above can be established in isolation and then community level response diversity calculated from individual species responses. However, this approach neglects any interspecific interactions within multispecies communities. Indeed, response diversity theory assumes that species-species relationships are less important for stability than abiotic environmental effects on species (that is, species-environment relationships; Ives & Carpenter 2007). Yet this assumption is not a limitation for two main reasons. First, one could simply model species responses (individually, using species-specific GAMs; Ross *et al*. 2022b) within a multispecies community, thereby allowing species-environment relationships to interact with species-species relationships as they would in nature (Ives & Carpenter 2007; Armitage & Jones 2020). Second, theoretical and empirical studies show that species-environment relationships are often more important than species-species relationships in driving both species asynchrony (*e*.*g*., Thibaut *et al*. 2012) and stability (*e*.*g*., Houlahan *et al*. 2007; Ives & Carpenter 2007). That interspecific interactions need not be included when measuring response diversity is therefore not necessarily a limiting assumption, but rather a realistic one. That said, future work should establish the relevance of different interaction types when empirically measuring response diversity (Windsor *et al*. 2023).

### 4.2 Multifarious environmental change

The framework above focuses on derivation of environment-dependent performance responses when a single environmental variable is varying. Yet, species and communities are often perturbed by multiple environmental stressors simultaneously. Rather than using response diversity solely as a predictor variable for estimating univariate stability responses, a system’s response along an environmental gradient could be measured as a function of a second environmental condition. That is, response diversity can be assessed for a given environmental state and compared to other treatments or replicates across any environmental ranges of interest, permitting quantification of response diversity to multiple environmental stressors, which often occur simultaneously (Dieleman *et al*. 2012; Bowler *et al*. 2020). Another approach, which we aim to develop, would be based on multivariate response surfaces (*e*.*g*., Yang *et al*. 2022). With this, multivariate response diversity may link conceptually to the study of co-tolerance to multiple stressors and stress-induced community sensitivity (see Vinebrooke *et al*. 2004) and should prove more operationalizable than univariate approaches given the propensity for multiple environmental stressors to not only co-occur but to interact nonadditively in nature (Dieleman *et al*. 2012). Similarly, multivariate response diversity metrics that simultaneously capture two or more performance responses to a given environmental driver could be developed. Where such responses involve several ecosystem functions, this should inform on the stability of ecosystem multifunctionality (Manning *et al*. 2018; Sasaki *et al*. 2019).

### 4.3 Abundance and scale-dependence

Response diversity depends on the identities and relative abundances of species within the community or functional group of interest. If community composition or abundance distributions change as a function of the environment, then response diversity should be abundance-weighted to account for species losses and gains along the environmental gradient—for example, by changing the parameter *q* involved in the calculation of the similarity-based metric used here (Leinster & Cobbold 2012). Such changes in species evenness can have knock-on consequences for species asynchrony (Thibaut & Connolly 2013), and stability (Hillebrand *et al*. 2008). Accounting for evenness in response diversity measures is therefore needed to capture how evenness effects scale from species responses to community or ecosystem stability. Moreover, species identities and abundances vary as a function of scale, making response diversity scale dependent. Practical users of response diversity should be careful not to extrapolate results beyond the focal temporal or spatial scale given that ecological stability and its drivers vary in time and space (Ross *et al*. 2021b; Jarillo *et al*. 2022). Disturbance also has a temporal and spatial signature, meaning that the spatiotemporal scale of reference determines the environmental conditions and potential disturbances to which ecological communities are subject (Zelnik *et al*. 2018; Jackson *et al*. 2021). Thus, response diversity measured over short timescales does not represent a holistic picture of how communities are likely to respond to the environment (that is, response diversity’s goal to mechanistically capture the insurance capacity of a system), since the range of environmental conditions experienced at shorter scales may be limited when compared to longer temporal windows (Wolkovich *et al*. 2014; Zelnik *et al*. 2018; Ross *et al*. 2021b). Despite the context-dependencies outlined here, response diversity has the potential to be a powerful tool for predicting ecological stability. Future studies should determine the temporal or spatial scales over which specific metrics of response diversity (*e*.*g*., dissimilarity, divergence, response trait diversity) perform best when aiming to predict ecological stability and whether the methods presented here require further refinement to better reflect the insurance capacity of ecological communities across scales.

### 4.4 Individual species contributions

Another extension of this framework would consider the relative importance of each species to community-level response diversity by, for example, measuring the median dissimilarity in performance-environment model slopes between each species and all others in the community. In this case, the species with the greatest dissimilarity from the others should be the largest contributor to similarity-based response diversity and divergence (*e*.*g*., *Colpidium striatum* in Figure 4d), and thus be the species whose loss from the community should most greatly reduce response diversity and erode stability (provided the two are mechanistically linked). Such an approach complements studies considering the contribution of individual species to ecological responses and stability (*e*.*g*., Donohue *et al*. 2017; White *et al*. 2020; Ross *et al*. 2021a, 2022a), in turn, aiding progress towards targeted conservation management decisions based on the quantifiable importance of individual taxa. Likewise, relative abundance and evenness are important when considering the contribution of species to response diversity, asynchrony, and stability (Thibaut & Connolly 2013). The similarity-based diversity metric we used here is based on hill numbers, with an initial value of *q* = 0 indicating a focus on species richness. By increasing *q* to higher values, it should be straightforward to also incorporate relative species abundances (Leinster & Cobbold 2012), and hence the potential role of dominance or rarity in shaping response diversity (*e*.*g*., Sasaki & Lauenroth 2011). Importantly, the subjects of response diversity analyses need not be different species; individuals can also exhibit diversity in their response traits, further underscoring the importance of intraspecific variation in ecological responses to the environment (Barabás & D’Andrea 2016; Mimura *et al*. 2017; Ross *et al*. 2017; Herrando-pérez *et al*. 2019).

### 4.5 Data resolution and spatiotemporal scope

Observational time series can also contain rich information on, for instance, community dynamics or environmental responses (Ives *et al*. 2003; Ushio *et al*. 2018; Shoemaker *et al*. 2022). Though methods for inferring causality from time series are emerging (*e*.*g*., *empirical dynamic modelling*, see Ushio *et al*. 2018; Ross *et al*. 2021b), such methods are often complex, and results must be interpreted carefully. The same is true if measuring response diversity from observational time series; though response diversity may be a key biotic mechanism underlying the insurance effect of biodiversity (Yachi & Loreau 1999), observational time series do not afford us information on causal relationships (but see Ushio *et al*. 2018). For dissimilarity, divergence, or other metrics of response diversity to be measurable from observational data, time series should comprise higher-level traits (*sensu* Agrawal *et al*. 2010) as described above (*e*.*g*., measuring intrinsic—that is, unlimited by competition—rather than realised rate of increase). Moreover, a high sampling resolution along environmental gradients is required to accurately differentiate between linear and nonlinear performance-environment relationships; longer or higher-resolution time series are better able to reveal nonlinear responses, and response diversity experiments should be designed with sampling resolution in mind to sufficiently capture all possible forms of performance-environment relationships. Compared to controlled experiments, observational data from the field is often ‘noisy’ and observation error can bias estimates of ecological interactions and response diversity (Ives *et al*. 2003), requiring additional care when measuring response diversity. As well as ensuring appropriate performance variables (*e*.*g*., vital rates) are measured that capture some aspect of species’ fitness or ecosystem functioning, one should aim to account for as many potentially confounding variables as possible when relating response diversity to stability, and to bear in mind that field studies rarely include all species from the regional pool, meaning response diversity is unlikely to portray the insurance capacity of the whole community. Future work should demonstrate the potential application of our response diversity framework to noisy field data. While the methods we describe here are useful for measuring stability from experimental or observational time series, dissimilarity and divergence could also be measured over spatial gradients to assess spatial variation in the stability and synchrony of functional properties such as primary productivity among habitat patches at a given spatial scale (Loreau *et al*. 2003; Lamy *et al*. 2019; Wang *et al*. 2019, 2021).

If response diversity can be measured from observational time series, it should be conceptually straightforward to harness global datasets (*e*.*g*., Dornelas *et al*. 2018) to measure response diversity at multiple sites across geographical regions, or to project locally-measured response diversity to other locations. Though our response diversity framework produces results that are necessarily a function of the environmental conditions against which species’ performances are assessed, comparing response diversity to relevant local environmental stressors across sites is informative insofar as it should predict stability. A global-scale analysis of response diversity to locally relevant environmental conditions should therefore unveil macroecological patterns and hotspots of ecological stability (Capdevila *et al*. 2022).

### 4.6 Environmental dependencies of response diversity and stability

That response diversity can be environment-dependent in cases where species respond nonlinearly to the environment has considerable implications for empirical studies of response diversity and for our understanding of ecological stability. Though threshold responses may be rare or difficult to detect in experiments (Hillebrand *et al*. 2020), nonlinear responses to the environment are common, and include functional responses to prey availability (Holling 1966; Daugaard *et al*. 2019), and physiological responses to light and temperature (*e*.*g*., Peek *et al*. 2002; Payne *et al*. 2016). Nonlinear responses to the environment may result in environment-dependencies in species asynchrony (Ghosh *et al*. 2020; Shoemaker *et al*. 2022). In turn, these can lead to environmentally dependent ecosystem multifunctionality and stability (Morin *et al*. 2014; Sasaki *et al*. 2019). Environment-dependent response diversity may therefore explain inconsistencies in the reported effects of biodiversity on ecological stability across studies (Jacquet *et al*. 2016; Pennekamp *et al*. 2018). Importantly, the empirical framework we propose permits identification of the specific environmental conditions under which response diversity should or should not be stabilising. For a nonlinear environmental response, *f*(*E*), one can examine the values of *E* (the ‘environment’) for which the derivatives of performance-environment relationships, *f*′(*E*), including their confidence regions, diverge (span zero). In other words, our framework makes it conceptually and practically simple to identify *a priori* hypotheses regarding the stabilising effect of response diversity based on environmental conditions. This should complement existing studies of the impact of environmental conditions on ecological stability (*e*.*g*., Hillebrand *et al*. 2018; Ross *et al*. 2022a) by identifying possible underlying biotic drivers of environmental responses. Moreover, in cases where response diversity varies as a function of the environment (that is, where performance-environment relationships are nonlinear), it is possible to manipulate response diversity without altering community composition. Comparing ecological stability between identical communities under different (fixed) environmental conditions, might then allow identification of response diversity’s mechanistic stabilising role. By controlling for species identity in this way, there are fewer potential confounding factors driving any observed community level responses (though environmental context is added as an extrinsic driver). For example, one could measure the diversity-stability relationship for a plant community with known performance responses to drought under drought and non-drought conditions to mechanistically consider how drought relates to stability. There is still much to learn about context-dependent ecological stability as it relates to nonlinear performance-environment relationships and environment-dependent response diversity, and we suggest this as a promising avenue of future work.

## 5. Conclusion

Since the observation that diversity alone is insufficient to explain the stabilising effect of species richness on aggregate ecosystem properties (May 1973; Ives *et al*. 1999), ecologists have been searching for a general mechanism explaining the insurance effect of biodiversity across contexts (Yachi & Loreau 1999). Here we proposed response diversity as a promising but understudied mechanism for predicting ecological responses to environmental conditions (Elmqvist *et al*. 2003; Nyström 2006; Mori *et al*. 2013). We found that empirical studies of response diversity were rare, and where they did occur, most measured the diversity of response traits (*e*.*g*., Laliberté *et al*. 2010; Morel *et al*. 2020). However, given the challenges associated with reliably producing response diversity estimates from low-level trait data (Bartomeus *et al*. 2017; Box 1), we suggest measuring response diversity more directly as the variation in performance responses to environmental conditions (Leary & Petchey 2009). The framework we proposed here provides a direct, empirically tractable, and conceptually simple estimate of response diversity based on dissimilarity and divergence in performance-environment relationships that should facilitate robust empirical tests of the diversity-stability relationship across taxa and environments. Such tests have the potential to identify the long sought-after mechanisms underpinning biodiversity’s role in buffering ecosystems against environmental change.

### Box 1.

#### Response trait diversity: a questionable indicator of stability?

Functional diversity measures the range and value of species’ traits which, in turn, aim to represent those species’ contributions ecosystem functioning (Tilman 2001). Myriad functional diversity metrics are now used to estimate ecosystem functioning using functional traits (Díaz & Cabido 2001; Petchey & Gaston 2006; Cadotte *et al*. 2011). Given their effect on ecosystem processes and functions, such traits are now termed *effect* traits, while *response* traits aim to capture information about how a species responds to environmental change (Suding *et al*. 2008; Oliver *et al*. 2015). Response traits are often assumed to, rather than empirically demonstrated to link to species’ environmental responses (Figure 1), and even where empirically tested, environmental responses are not always well captured by putative functional traits (Bartomeus *et al*. 2017). Examples of response traits include life-form and seed mass in plants (Sasaki *et al*. 2019), aquatic stage and reproductive strategy in freshwater macroinvertebrates (Thornhill *et al*. 2018), and clutch size and thermal maximum in birds (Hordley *et al*. 2021). Response trait diversity is then typically measured as the functional dispersion (FDis) of the response trait space (Laliberté *et al*. 2010; Rader *et al*. 2013), or sometimes using the closely related Rao’s Quadratic Entropy (RaoQ) index (Chillo *et al*. 2011; Correia *et al*. 2018). Some studies alternatively group species into response groups (*e*.*g*., Joseph *et al*. 2014; Schweiger *et al*. 2015), where species within a functional group are posited to respond similarly to environmental change based on their similar response traits. Studies aiming to quantify functional redundancy have therefore typically relied on functional grouping (*e*.*g*., Laliberté *et al*. 2010; Bruno *et al*. 2016). However, this approach ignores a suite of functional differences between species; functional grouping is rarely ecologically meaningful (Petchey & Gaston 2006).

Considerable challenges exist when quantifying functional diversity from traits and linking traits to ecological stability. For example, there is still no consensus on which traits are ‘functional’ in a range of systems (Violle *et al*. 2007), and many traits can be highly correlated with, for example, body size (Kozłowski & Weiner 1997). When attempting to measure response trait diversity, it can be difficult to delineate responses from effects; many traits, such as specific leaf area or photosynthetic pathway (Sasaki *et al*. 2019), act both as response and effect traits depending on environmental context (Hooper *et al*. 2005; Suding *et al*. 2008). This difficulty in separating response and effect traits could be a critical flaw in many trait diversity studies given that the overlap between the identity of response and effect traits is expected to drive changes in ecosystem functioning with the environment (Suding & Goldstein 2008); as the overlap between response and effect traits increases, changes in the environment more likely modify ecosystem functioning. Accordingly, ecosystem functioning is expected to be stable only in cases where effect traits and response traits do not overlap, else changes in response diversity will necessarily propagate to affect ecosystem functioning (de Bello *et al*. 2021).

It is also often practically challenging to measure functional traits in the field, particularly for mobile organisms (Pakeman & Quested 2007; Moretti *et al*. 2017). When measuring functional diversity from species traits, there is no single best metric for all scenarios, as most functional diversity indices capture distinct aspects of the functional differences within a community (Villéger *et al*. 2008; Schleuter *et al*. 2010). Linking trait diversity and stability through species asynchrony requires examining multiple response traits simultaneously (van Klink *et al*. 2019), which involves considerable sampling effort, as well as categorisation of traits as response and/or effect traits.

Groups of species with similar response trait values should exhibit more synchronous population dynamics, while those with more distinct response traits are more likely to fluctuate asynchronously (Adler *et al*. 2013; Lepš *et al*. 2018). As communities are exposed to a wider range of environmental conditions, the importance of multi-trait dissimilarity in driving species asynchrony increases. Conversely, when interested in a specific environmental perturbation, individual response trait values may be more important for species’ performance and ecosystem functioning than multidimensional response trait diversity (de Bello *et al*. 2021). Another metric of interest may be Community-Weighted Mean (CWM) trait values, particularly when considering the effect of dominant species and their traits on stability (Sasaki & Lauenroth 2011; de Bello *et al*. 2021).

Further methodological considerations include whether and how to abundance-weight functional diversity indices to account for rare species (Lavorel *et al*. 2008), and how to handle intraspecific trait variation given its ecological importance (de Bello *et al*. 2011; Ross *et al*. 2017; Herrando-pérez *et al*. 2019). Another important consideration if linking trait diversity and stability is *when* to measure response trait diversity. Most studies linking traits and stability consider the response trait diversity of some average community through time (*e*.*g*., Craven *et al*. 2018), but this ignores temporal community turnover, thereby weakening the expected relationship between response trait diversity and stability (de Bello *et al*. 2021). Together, these challenges and others considerably limit the efficacy of using low-level functional traits to empirically quantify response diversity and measure its relationship with temporal stability. Unless using well-defined and empirically tested response traits (*e*.*g*., Kandlikar *et al*. 2022), we therefore suggest caution in measuring response diversity from low-level functional traits, and in assuming that such trait diversity should reflect any meaningful facet of ecological stability.

## Acknowledgements

We thank Dan Leary for generating the empirical data we reanalysed in this study, Helmut Hillebrand, Charlotte Kunze, and Frank Pennekamp for helpful discussion, and five anonymous reviewers for constructive comments on earlier drafts. This study was supported by a SHINKA collaboration grant awarded to SRP-JR by the Okinawa Institute of Science and Technology Graduate University (OIST). SRP-JR and DWA were supported by subsidy funding to OIST. OLP was supported by the University of Zurich Research Priority Programme in Global Change and Biodiversity (URPP GCB) and by SNF project 310030_188431. TS was supported by a Fostering Joint International Research A (no. 19KK0393) from the Japanese Ministry of Education, Culture, Sports, Science and Technology.

## Data availability statement

All data and R code necessary to reproduce the analyses and figures are available in the Zenodo digital repository https://doi.org/10.5281/zenodo.7018707 (Ross *et al*. 2022b).

## Conflict of ihypernterest

The authors declare no conflicts of interest.

## References

Adler, P.B., Fajardo, A., Kleinhesselink, A.R., & Kraft, N.J. (2013). Trait-based tests of coexistence mechanisms. Ecol. Lett., 16, 1294–1306.

Agrawal, A.A., Conner, J.K., & Rasmann, S. (2010). Tradeoffs and negative correlations in evolutionary ecology. Evolution since Darwin: the first 150 years. pg. 243–268.

Allen C.R., Cumming G.S., Garmestani A.S., Taylor P.D., Walker B.H. (2011). Managing for resilience. Wildlife Biology, 17, 337–349.

Altomare, M., Vasconcelos, H.L., Raymundo, D., Lopes, S., Vale, V. & Prado-Junior, J. (2021). Assessing the fire resilience of the savanna tree component through a functional approach. Acta Oecologica, 111, 103728.

Armitage, D.W. & Jones, S.E. (2019). Negative frequency-dependent growth underlies the stable coexistence of two cosmopolitan aquatic plants. Ecology, 100, e02657.

Armitage, D.W. & Jones, S.E. (2020). Coexistence barriers confine the poleward range of a globally distributed plant. Ecol. Lett., 23, 1838–1848.

Arnoldi, J.F., Loreau, M. & Haegeman, B. (2019). The inherent multidimensionality of temporal variability: how common and rare species shape stability patterns. Ecol. Lett., 22, 1557–1567.

Barabás, G. & D’Andrea, R. (2016). The effect of intraspecific variation and heritability on community pattern and robustness. Ecol. Lett., 19, 977–986.

Bartomeus, I., Cariveau, D.P., Harrison, T. & Winfree, R. (2017). On the inconsistency of pollinator species traits for predicting either response to land-use change or functional contribution. Oikos, 127, 306–315.

Bartomeus, I., Park, M.G., Gibbs, J., Danforth, B.N., Lakso, A.N. & Winfree, R. (2013). Biodiversity ensures plant– pollinator phenological synchrony against climate change. Ecol. Lett., 16, 1331–1338.

Baskett, M.L., Fabina, N.S. & Gross, K. (2014). Response diversity can increase ecological resilience to disturbance in coral reefs. Am. Nat., 184, E16–E31.

de Bello, F., Lavorel, S., Albert, C.H., Thuiller, W., Grigulis, K., Dolezal, J., et al. (2011). Quantifying the relevance of intraspecific trait variability for functional diversity. Methods Ecol. Evol., 2, 163–174.

de Bello, F., Lavorel, S., Hallett, L.M., Valencia, E., Garnier, E., Roscher, C., et al. (2021). Functional trait effects on ecosystem stability: assembling the jigsaw puzzle. Trends Eco. Evo., 36, 822–836.

Berdugo, M., Delgado-Baquerizo, M., Soliveres, S., Hernández-Clemente, R., Zhao, Y., Gaitán, J.J., et al. (2020). Global ecosystem thresholds driven by aridity. Science, 367, 787–790.

Bowler, D.E., Bjorkman, A.D., Dornelas, M., Myers-Smith, I.H., Navarro, L.M., Niamir, A., et al. (2020). Mapping human pressures across the planet uncovers anthropogenic threat complexes. People Nat., 2, 380– 394.

Bruno, D., Gutiérrez-Cánovas, C., Sánchez-Fernández, D., Velasco, J., & Nilsson, C. (2016). Impacts of environmental filters on functional redundancy in riparian vegetation. J. Appl. Ecol., 53, 846–855.

Cadotte, M.W., Carscadden, K. & Mirotchnick, N. (2011). Beyond species: Functional diversity and the maintenance of ecological processes and services. J. Appl. Ecol., 48, 1079–1087.

Capdevila, P., Noviello, N., McRae, L., Freeman, R. & Clements, C.F. (2022). Global patterns of resilience decline in vertebrate populations. Ecol. Lett., 25, 240–251.

Cariveau, D.P., Williams, N.M., Benjamin, F.E. & Winfree, R. (2013). Response diversity to land use occurs but does not consistently stabilise ecosystem services provided by native pollinators. Ecol. Lett., 16, 903– 911.

Chesson, P. (2000). Mechanisms of Maintenance of Species Diversity. Annu. Rev. Ecol. Syst., 31, 343–366.

Chillo, V., Anand, M. & Ojeda, R.A. (2011). Assessing the Use of Functional Diversity as a Measure of Ecological Resilience in Arid Rangelands. Ecosystems, 14, 1168–1177.

Correia, D.L.P., Raulier, F., Bouchard, M. & Filotas, É. (2018). Response diversity, functional redundancy, and post-logging productivity in northern temperate and boreal forests. Ecol. Appl., 28, 1282–1291.

Craven, D., Eisenhauer, N., Pearse, W.D., Hautier, Y., Isbell, F., Roscher, C., et al. (2018). Multiple facets of biodiversity drive the diversity–stability relationship. Nat. Ecol. Evol., 2, 1579–1587.

Craven, D., Filotas, E., Angers, V.A. & Messier, C. (2016). Evaluating resilience of tree communities in fragmented landscapes: linking functional response diversity with landscape connectivity. Divers. Distrib., 22, 505– 518.

Daugaard, U., Petchey, O.L. & Pennekamp, F. (2019). Warming can destabilize predator–prey interactions by shifting the functional response from Type III to Type II. J. Anim. Ecol., 88, 1575–1586.

Díaz, S. & Cabido, M. (2001). Vive la différence: Plant functional diversity matters to ecosystem processes. Trends Ecol. Evol., 16, 646–655.

Dieleman, W.I., Vicca, S., Dijkstra, F.A., Hagedorn, F., Hovenden, M.J., Larsen, K.S., et al. (2012). Simple additive effects are rare: a quantitative review of plant biomass and soil process responses to combined manipulations of CO2 and temperature. Glob. Chang. Biol., 18, 2681–2693.

Döbert, T.F., Webber, B.L., Sugau, J.B., Dickinson, K.J.M. & Didham, R.K. (2017). Logging increases the functional and phylogenetic dispersion of understorey plant communities in tropical lowland rain forest. J. Ecol., 105, 1235–1245.

Donohue, I., Petchey, O.L., Kéfi, S., Génin, A., Jackson, A.L., Yang, Q., et al. (2017). Loss of predator species, not intermediate consumers, triggers rapid and dramatic extinction cascades. Glob. Change Biol., 23, 2962– 2972.

Donohue, I., Petchey, O.L., Montoya, J.M., Jackson, A.L., Mcnally, L., Viana, M., et al. (2013). On the dimensionality of ecological stability. Ecol. Lett., 16, 421–429.

Dornelas, M., Antão, L.H., Moyes, F., Bates, A.E., Magurran, A.E., Adam, D., et al. (2018). BioTIME: A database of biodiversity time series for the Anthropocene. Glob. Ecol. Biogeogr., 27, 760–786.

Elmqvist, T., Folke, C., Nyström, M., Peterson, G., Bengtsson, J., Walker, B., et al. (2003). Response diversity, ecosystem change, and resilience. Front. Ecol. Environ., 1, 488–494.

Elton, C.S. (1958). Ecology of Invasions by Animals and Plants. Chapman & Hall, London.

Fauchald, P., Skov, H., Skern-Mauritzen, M., Hausner, V.H., Johns, D. & Tveraa, T. (2011). Scale-dependent response diversity of seabirds to prey in the North Sea. Ecology, 92, 228–239.

Ford, H., Healey, J.R., Markesteijn, L. & Smith, A.R. (2018). How does grazing management influence the functional diversity of oak woodland ecosystems? A plant trait approach. Agric. Ecosyst. Environ., 258, 154–161.

Fründ, J., Zieger, S.L. & Tscharntke, T. (2013). Response diversity of wild bees to overwintering temperatures. Oecologia, 173, 1639–1648.

Ghosh, S., Sheppard, L.W., Reid, P.C. & Reuman, D. (2020). A new approach to interspecific synchrony in population ecology using tail association. Ecol. Evol., 10, 12764–12776.

Gonzalez, A. & Loreau, M. (2009). The Causes and Consequences of Compensatory Dynamics in Ecological Communities. Annu. Rev. Ecol. Evol. Syst., 40, 393–414.

Grainger, T.N., Levine, J.M. & Gilbert, B. (2019). The Invasion Criterion: A Common Currency for Ecological Research. Trends Ecol. Evol., 34, 925–935.

Hammill, E., Kratina, P., Vos, M., Petchey, O.L. & Anholt, B.R. (2015). Food web persistence is enhanced by non-trophic interactions. Oecologia, 178, 549–556.

Hastie, T. & Tibshirani, R. (1987). Generalized additive models: some applications. J. Am. Stat. Assoc., 82, 371– 386.

Hector, A.A., Hautier, Y., Saner, P., Wacker, L., Bagchi, R., Joshi, J., et al. (2015). General stabilizing effects of plant diversity on grassland productivity through population asynchrony and overyielding. Ecology, 91, 2213–2220.

Herrando-pérez, S., Vieites, D.R., Monasterio, F.F.C., Belliure, J., Chown, S.L., Beukema, W., et al. (2019). Intraspecific variation in lizard heat tolerance alters estimates of climate impact. J. Anim. Ecol., 88, 247–257.

Hillebrand, H., Bennett, D.M., & Cadotte, M.W. (2008). Consequences of dominance: a review of evenness effects on local and regional ecosystem processes. Ecology, 89, 1510–1520.

Hillebrand, H., Donohue, I., Harpole, W.S., Hodapp, D., Kucera, M., Lewandowska, A.M., et al. (2020). Thresholds for ecological responses to global change do not emerge from empirical data. Nat. Ecol. Evol., 4, 1502– 1509.

Hillebrand, H., Langenheder, S., Lebret, K., Lindström, E., Östman, Ö. & Striebel, M. (2018). Decomposing multiple dimensions of stability in global change experiments. Ecol. Lett., 21, 21–30.

Holling, C.S. (1966). The Functional Response of Invertebrate Predators to Prey Density. Mem. Entomol. Soc. Can., 98, 5–86.

Hooper, D.U., Chapin, F.S., Ewel, J.J., Hector, A., Inchausti, P., Lavorel, S., et al. (2005). Effects of biodiversity on ecosystem functioning: A consensus of current knowledge. Ecol. Monogr., 75, 3–35.

Hordley, L.A., Gillings, S., Petchey, O.L., Tobias, J.A. & Oliver, T.H. (2021). Diversity of response and effect traits provides complementary information about avian community dynamics linked to ecological function. Funct. Ecol., 35, 1938–1950.

Houlahan, J.E., Currie, D.J., Cottenie, K., Cumming, G.S., Ernest, S.K.M., Findlay, C.S., et al. (2007). Compensatory dynamics are rare in natural ecological communities. Proc. Natl. Acad. Sci. USA, 104, 3273–3277.

Ishii, R. & Masahiko, M. (2001). Coexistence induced by pollen limitation in flowering-plant species. Proc. R. Soc. Lond. B Biol. Sci., 268, 579–585.

Ives, A.R. & Carpenter, S.R. (2007). Stability and diversity of ecosystems. Science, 317, 58–62.

Ives, A.R., Dennis, B., Cottingham, K.L. & Carpenter, S.R. (2003). Estimating community stability and ecological interactions from time-series data. Ecol. monogr., 73, 301–330.

Ives, A.R., Gross, K. & Klug, J.L. (1999). Stability and Variability in Competitive Communities. Science, 286, 542– 544.

Jackson, M.C., Pawar, S., & Woodward, G. (2021). The temporal dynamics of multiple stressor effects: from individuals to ecosystems. Trends Eco. Evo., 36, 402–410.

Jacquet, C., Moritz, C., Morissette, L., Legagneux, P., Massol, F., Archambault, P., et al. (2016). No complexity– stability relationship in empirical ecosystems. Nat. Commun., 7, 12573.

Jarillo, J., Cao-García, F.J. & De Laender, F. (2022). Spatial and ecological scaling of stability in spatial community networks. Front. Ecol. Evol., 10.

Joseph, G.S., Seymour, C.L., Cumming, G.S., Cumming, D.H.M. & Mahlangu, Z. (2014). Termite Mounds Increase Functional Diversity of Woody Plants in African Savannas. Ecosystems, 17, 808–819.

Kahiluoto, H., Kaseva, J., Olesen, J.E., Kersebaum, K.C., Ruiz-Ramos, M., Gobin, A., et al. (2019). Genetic response diversity to provide yield stability of cultivar groups deserves attention. Proc. Natl. Acad. Sci. U.S.A., 166, 10627–10629.

Kandlikar, G.S., Kleinhesselink, A.R. & Kraft, N.J.B. (2022). Functional traits predict species responses to environmental variation in a California grassland annual plant community. J. Ecol. [In press].

Kondoh, M. (2003). Foraging Adaptation and the Relationship Between Food-Web Complexity and Stability. Science, 299, 1388–1391.

Kozłowski, J. & Weiner, J. (1997). Interspecific allometries Are by-products of body size optimization. Am. Nat., 149, 352–380.

Laliberté, E., Wells, J.A., Declerck, F., Metcalfe, D.J., Catterall, C.P., Queiroz, C., et al. (2010). Land-use intensification reduces functional redundancy and response diversity in plant communities. Ecol. Lett., 13, 76–86.

Lamy, T., Wang, S., Renard, D., Lafferty, K.D., Reed, D.C. & Miller, R.J. (2019). Species insurance trumps spatial insurance in stabilizing biomass of a marine macroalgal metacommunity. Ecology, 100, e02719.

Lavorel, S., Grigulis, K., McIntyre, S., Williams, N.S.G., Garden, D., Dorrough, J., et al. (2008). Assessing functional diversity in the field -Methodology matters! Funct. Ecol., 22, 134–147.

Leary, D.J. & Petchey, O.L. (2009). Testing a biological mechanism of the insurance hypothesis in experimental aquatic communities. J. Anim. Ecol., 78, 1143–1151.

Leinster, T. & Cobbold, C.A. (2012). Measuring diversity: the importance of species similarity. Ecology, 93, 477– 489.

Lepš, J., Májeková, M., Vítová, A., Doležal, J., & de Bello, F. (2018). Stabilizing effects in temporal fluctuations: management, traits, and species richness in high-diversity communities. Ecology, 99, 360–371.

Loreau, M. (2010). Stability and Complexity of Ecosystems: New Perspectives on an Old Debate. In: From Populations to Ecosystems, Theoretical Foundations for a New Ecological Synthesis (MPB-46). Princeton University Press, pp. 123–163.

Loreau, M., Barbier, M., Filotas, E., Gravel, D., Isbell, F., Miller, S.J., et al. (2021). Biodiversity as insurance: from concept to measurement and application. Biol. Rev., 96, 2333–2354.

Loreau, M. & de Mazancourt, C. (2013). Biodiversity and ecosystem stability: A synthesis of underlying mechanisms. Ecol. Lett., 16, 106–115.

Loreau, M., Mouquet, N., Gonzalez, A. & Mooney, H.A. (2003). Biodiversity as spatial insurance in heterogeneous landscapes. Proc. Natl. Acad. Sci. U.S.A., 100, 12765–12770.

MacArthur, R. (1955). Fluctuations of Animal Populations and a Measure of Community Stability. Ecology, 36, 533–536.

Malyshev, A.V., Arfin Khan, M.A.S., Beierkuhnlein, C., Steinbauer, M.J., Henry, H.A.L., Jentsch, A., et al. (2016). Plant responses to climatic extremes: within-species variation equals among-species variation. Glob. Change Biol., 22, 449–464.

Mandle, L. & Ticktin, T. (2015). Moderate land use changes plant functional composition without loss of functional diversity in India’s Western Ghats. Ecol. Appl., 25, 1711–1724.

Manning, P., Van Der Plas, F., Soliveres, S., Allan, E., Maestre, F.T., Mace, G., et al. (2018). Redefining ecosystem multifunctionality. Nat. Ecol. Evol. 2, 427–436.

May, R.M. (1973). Stability and complexity in model ecosystems. Princeton University Press.

McCann, K., Hastings, A. & Huxel, G.R. (1998). Weak trophic interactions and the balance of nature. Nat. 1998 3956704, 395, 794–798.

McCann, K.S. (2000). The diversity-stability debate. Nature, 405, 228–233.

McCann, M.J. (2016). Response diversity of free-floating plants to nutrient stoichiometry and temperature: growth and resting body formation. PeerJ, 4, e1781.

Mimura, M., Yahara, T., Faith, D.P., Vázquez-Domínguez, E., Colautti, R.I., Araki, H., et al. (2017). Understanding and monitoring the consequences of human impacts on intraspecific variation. Evol. Appl., 10, 121– 139.

Moore, J.W. & Olden, J.D. (2017). Response diversity, nonnative species, and disassembly rules buffer freshwater ecosystem processes from anthropogenic change. Glob. Change Biol., 23, 1871–1880.

Morel, L., Barbe, L., Jung, V., Clément, B., Schnitzler, A. & Ysnel, F. (2020). Passive rewilding may (also) restore phylogenetically rich and functionally resilient forest plant communities. Ecol. Appl., 30, e02007.

Moretti, M., Dias, A.T.C., de Bello, F., Altermatt, F., Chown, S.L., Azcárate, F.M., et al. (2017). Handbook of protocols for standardized measurement of terrestrial invertebrate functional traits. Funct. Ecol., 31, 558–567.

Mori, A.S., Furukawa, T. & Sasaki, T. (2013). Response diversity determines the resilience of ecosystems to environmental change. Biol. Rev., 88, 349–364.

Morin, X., Fahse, L., de Mazancourt, C., Scherer-Lorenzen, M. & Bugmann, H. (2014). Temporal stability in forest productivity increases with tree diversity due to asynchrony in species dynamics. Ecol. Lett., 17, 1526– 1535.

Mougi, A. & Kondoh, M. (2012). Diversity of Interaction Types and Ecological Community Stability. Science, 337, 349–351.

Mumme, S., Jochum, M., Brose, U., Haneda, N.F. & Barnes, A.D. (2015). Functional diversity and stability of litter-invertebrate communities following land-use change in Sumatra, Indonesia. Biol. Conserv., 191, 750–758.

Nyström, M. (2006). Redundancy and Response Diversity of Functional Groups: Implications for the Resilience of Coral Reefs. AMBIO J. Hum. Environ., 35, 30–35.

Odum, E.P. (1953). Fundamentals of Ecology. Saunders, Philadelphia.

Oliver, T.H., Heard, M.S., Isaac, N.J.B., Roy, D.B., Procter, D., Eigenbrod, F., et al. (2015). Biodiversity and Resilience of Ecosystem Functions. Trends Ecol. Evol., 30, 673–684.

Pakeman, R.J. & Quested, H.M. (2007). Sampling plant functional traits: What proportion of the species need to be measured? Appl. Veg. Sci., 10, 91–96.

Payne, N.L., Smith, J.A., van der Meulen, D.E., Taylor, M.D., Watanabe, Y.Y., Takahashi, A., et al. (2016). Temperature dependence of fish performance in the wild: links with species biogeography and physiological thermal tolerance. Funct. Ecol., 30, 903–912.

Peek, M.S., Russek-Cohen, E., Wait, A.D. & Forseth, I.N. (2002). Physiological response curve analysis using nonlinear mixed models. Oecologia, 132, 175–180.

Pennekamp, F., Pontarp, M., Tabi, A., Altermatt, F., Alther, R., Choffat, Y., et al. (2018). Biodiversity increases and decreases ecosystem stability. Nature, 563, 109–112.

Petchey, O.L. (2003). Integrating methods that investigate how complementarity influences ecosystem functioning. Oikos, 101, 323–330.

Petchey, O.L. & Gaston, K.J. (2006). Functional diversity: Back to basics and looking forward. Ecol. Lett., 9, 741– 758.

Pimm, S.L. (1984). The complexity and stability of ecosystems. Nature, 307, 321–326.

Pimm, S.L. & Lawton, J.H. (1978). On feeding on more than one trophic level. Nature, 275, 542–544.

Pinek, L., Mansour, I., Lakovic, M., Ryo, M., & Rillig, M.C. (2020). Rate of environmental change across scales in ecology. Biol. Rev., 95, 1798–1811.

Rader, R., Reilly, J., Bartomeus, I. & Winfree, R. (2013). Native bees buffer the negative impact of climate warming on honey bee pollination of watermelon crops. Glob. Change Biol., 19, 3103–3110.

Reilly, J.R., Artz, D.R., Biddinger, D., Bobiwash, K., Boyle, N.K., Brittain, C., et al. (2020). Crop production in the USA is frequently limited by a lack of pollinators. Proc. R. Soc. B Biol. Sci., 287, 20200922.

Ross, S.R.P.-J., Arnoldi, J.-F., Loreau, M., White, C.D., Stout, J.C., Jackson, A.L., et al. (2021a). Universal scaling of robustness of ecosystem services to species loss. Nat. Commun., 12, 5167.

Ross, S.R.P.-J., García Molinos, J., Okuda, A., Johnstone, J., Atsumi, K., Futamura, R., et al. (2022a). Predators mitigate the destabilising effects of heatwaves on multitrophic stream communities. Glob. Change Biol., 28, 403–416.

Ross, S.R.P.-J., Hassall, C., Hoppitt, W.J.E., Edwards, F.A., Edwards, D.P. & Hamer, K.C. (2017). Incorporating intraspecific trait variation into functional diversity: Impacts of selective logging on birds in Borneo. Methods Ecol. Evol., 8, 1499–1505.

Ross, S.R.P.-J., Petchey, O.L., Sasaki, T., & Armitage, D.W. (2022b). Supporting Information: opetchey/response_diversity_how_to_measure: v1.1-review (Pre-release). Zenodo. DOI: 10.5281/zenodo.7018707

Ross, S.R.P.-J., Suzuki, Y., Kondoh, M., Suzuki, K., Villa Martín, P. & Dornelas, M. (2021b). Illuminating the intrinsic and extrinsic drivers of ecological stability across scales. Ecol. Res., 36, 364–378.

Sasaki, T. & Lauenroth, W.K. (2011). Dominant species, rather than diversity, regulates temporal stability of plant communities. Oecologia, 166, 761–768.

Sasaki, T., Lu, X., Hirota, M. & Bai, Y. (2019). Species asynchrony and response diversity determine multifunctional stability of natural grasslands. J. Ecol., 107, 1862–1875.

Schleuter, D., Daufresne, M., Massol, F. & Argillier, C. (2010). A user’s guide to functional diversity indices. Ecol. Monogr., 80, 469–484.

Schnabel, F., Liu, X., Kunz, M., Barry, K.E., Bongers, F.J., Bruelheide, H., et al. (2021). Species richness stabilizes productivity via asynchrony and drought-tolerance diversity in a large-scale tree biodiversity experiment. Sci. Adv., 7, eabk1643.

Schweiger, A.H., Audorff, V. & Beierkuhnlein, C. (2015). The acid taste of climate change: 20th century acidification is re-emerging during a climatic extreme event. Ecosphere, 6, art94.

Shade, A. (2017). Diversity is the question, not the answer. ISME J., 11, 1–6.

Shoemaker, L.G., Hallett, L.M., Zhao, L., Reuman, D.C., Wang, S., Cottingham, K.L., et al. (2022). The long and the short of it: Mechanisms of synchronous and compensatory dynamics across temporal scales. Ecology, e3650.

Shyu, E., & Caswell, H. (2014). Calculating second derivatives of population growth rates for ecology and evolution. Methods Ecol. Evol., 5, 473–482.

Simpson, G.L. (2021). gratia: Graceful ‘ggplot’-Based Graphics and Other Functions for GAMs Fitted Using ‘mgcv’. R package version 0.6.0. https://CRAN.R-project.org/package=gratia

Spasojevic, M.J., Bahlai, C.A., Bradley, B.A., Butterfield, B.J., Tuanmu, M.-N., Sistla, S., et al. (2016). Scaling up the diversity–resilience relationship with trait databases and remote sensing data: the recovery of productivity after wildfire. Glob. Change Biol., 22, 1421–1432.

Stavert, J.R., Pattemore, D.E., Gaskett, A.C., Beggs, J.R. & Bartomeus, I. (2017). Exotic species enhance response diversity to land-use change but modify functional composition. Proc. R. Soc. B Biol. Sci., 284, 20170788.

Suding, K.N., & Goldstein, L.J. (2008). Testing the Holy Grail framework: using functional traits to predict ecosystem change. New Phytol., 180, 559–562.

Suding, K.N., Lavorel, S., Chapin, F.S., Cornelissen, J.H.C., Díaz, S., Garnier, E., et al. (2008). Scaling environmental change through the community-level: A trait-based response-and-effect framework for plants. Glob. Change Biol., 14, 1125–1140.

Thebault, E. & Fontaine, C. (2010). Stability of Ecological Communities and the Architecture of Mutualistic and Trophic Networks. Science, 329, 853–856.

Thibaut, L.M., & Connolly, S.R. (2013) Understanding diversity–stability relationships: towards a unified model of portfolio effects. Ecol. Lett., 16, 140–150.

Thibaut, L.M., Connolly, S.R. & Sweatman, H.P.A. (2012). Diversity and stability of herbivorous fishes on coral reefs. Ecology, 93, 891–901.

Thornhill, I., Biggs, J., Hill, M., Briers, R., Gledhill, D., Wood, P., et al. (2018). The functional response and resilience in small waterbodies along land-use and environmental gradients. Glob. Change Biol., 24, 3079–3092.

Tilman, D. (2001). Functional Diversity. In: Encyclopedia of Biodiversity. Elsevier Inc., pp. 587–596.

Tilman, D., Lehman, C.L. & Bristow, C.E. (1998). Diversity-stability relationships: statistical inevitability or ecological consequence? Am. Nat., 151, 277–82.

Ushio, M., Hsieh, C.H., Masuda, R., Deyle, E.R., Ye, H., Chang, C.W., et al. (2018). Fluctuating interaction network and time-varying stability of a natural fish community. Nature, 554, 360–363.

van Klink, R., Lepš, J., Vermeulen, R., & de Bello, F. (2019). Functional differences stabilize beetle communities by weakening interspecific temporal synchrony. Ecology, 100, e02748.

Villa Martín, P., Muñoz, M.A., & Pigolotti, S. (2019). Bet-hedging strategies in expanding populations. PLoS Comp. Biol., 15, e1006529.

Villéger, S., Mason, N.W.H. & Mouillot, D. (2008). New multidimensional functional diversity indices for a multifaceted framwork in functional ecology. Ecology, 89, 2290–2301.

Vinebrooke, R.D., Cottingham, K.L., Norberg, J., Scheffer, M., Dodson, S.I., Maberly, S.C., et al. (2004). Impacts of multiple stressors on biodiversity and ecosystem functioning: The role of species co-tolerance. Oikos, 104, 451–457.

Violle, C., Navas, M.L., Vile, D., Kazakou, E., Fortunel, C., Hummel, I., et al. (2007). Let the concept of trait be functional! Oikos, 116, 882–892.

Vogel, A., Manning, P., Cadotte, M.W., Cowles, J., Isbell, F., Jousset, A.L.C., et al. (2019). Lost in trait space: species-poor communities are inflexible in properties that drive ecosystem functioning. Adv. Ecol. Res. 1st edn. Elsevier Ltd.

Walker, B., Kinzig, A. & Langridge, J. (1999). Plant Attribute Diversity, Resilience, and Ecosystem Function: The Nature and Significance of Dominant and Minor Species. Ecosystems, 2, 95–113.

Wang, S. & Loreau, M. (2016). Biodiversity and ecosystem stability across scales in metacommunities. Ecol. Lett., 19, 510–518.

Wang, S., Loreau, M., de Mazancourt, C., Isbell, F., Beierkuhnlein, C., Connolly, J., et al. (2021). Biotic homogenization destabilizes ecosystem functioning by decreasing spatial asynchrony. Ecology, 102, 1– 10.

Wang, Y., Cadotte, M.W., Chen, Y., Fraser, L.H., Zhang, Y., Huang, F., et al. (2019). Global evidence of positive biodiversity effects on spatial ecosystem stability in natural grasslands. Nat. Commun., 10, 3207–3207.

White, L., O’Connor, N.E., Yang, Q., Emmerson, M.C. & Donohue, I. (2020). Individual species provide multifaceted contributions to the stability of ecosystems. Nat. Ecol. Evol., 4, 1594–1601.

Windsor, F.M., van den Hoogen, J., Crowther, T.W., & Evans, D.M. (2023). Using ecological networks to answer questions in global biogeography and ecology. J. Biogeog., 50, 57–69.

Winfree, R. & Kremen, C. (2009). Are ecosystem services stabilized by differences among species? A test using crop pollination. Proc. R. Soc. B Biol. Sci., 276, 229–237.

Wolkovich, E.M., Cook, B.I., McLauchlan, K.K., & Davies, T.J. (2014). Temporal ecology in the Anthropocene. Ecol. Lett., 17, 1365–1379.

Wood, S.N., Pya, N. & Säfken, B. (2016). Smoothing Parameter and Model Selection for General Smooth Models. J. Am. Stat. Assoc., 111, 1548–1563.

Yachi, S. & Loreau, M. (1999). Biodiversity and ecosystem productivity in a fluctuating environment: The insurance hypothesis. Proc. Natl. Acad. Sci., 96, 1463–1468.

Yang, G., Ryo, M., Roy, J., Lammel, D.R., Ballhausen, M-B., Jing, X., et al. (2022). Multiple anthropogenic pressures eliminate the effects of soil microbial diversity on ecosystem functions in experimental microcosms. Nat. Commun., 13, 4260.

Yodzis, P. (1981). The stability of real ecosystems. Nature, 289, 674–676.

Zelnik, Y., Arnoldi, J.-F. & Loreau, M. (2018). The impact of spatial and temporal dimensions of disturbances on ecosystem stability. Front. Ecol. Evol., 6, 224.

