## Supplementary Information for "How to measure response diversity"

Contents

Pg. 2. **Figure S1: PRISMA diagram.**

Pg 3-5. **List of papers included in final results after screening for eligibility**.

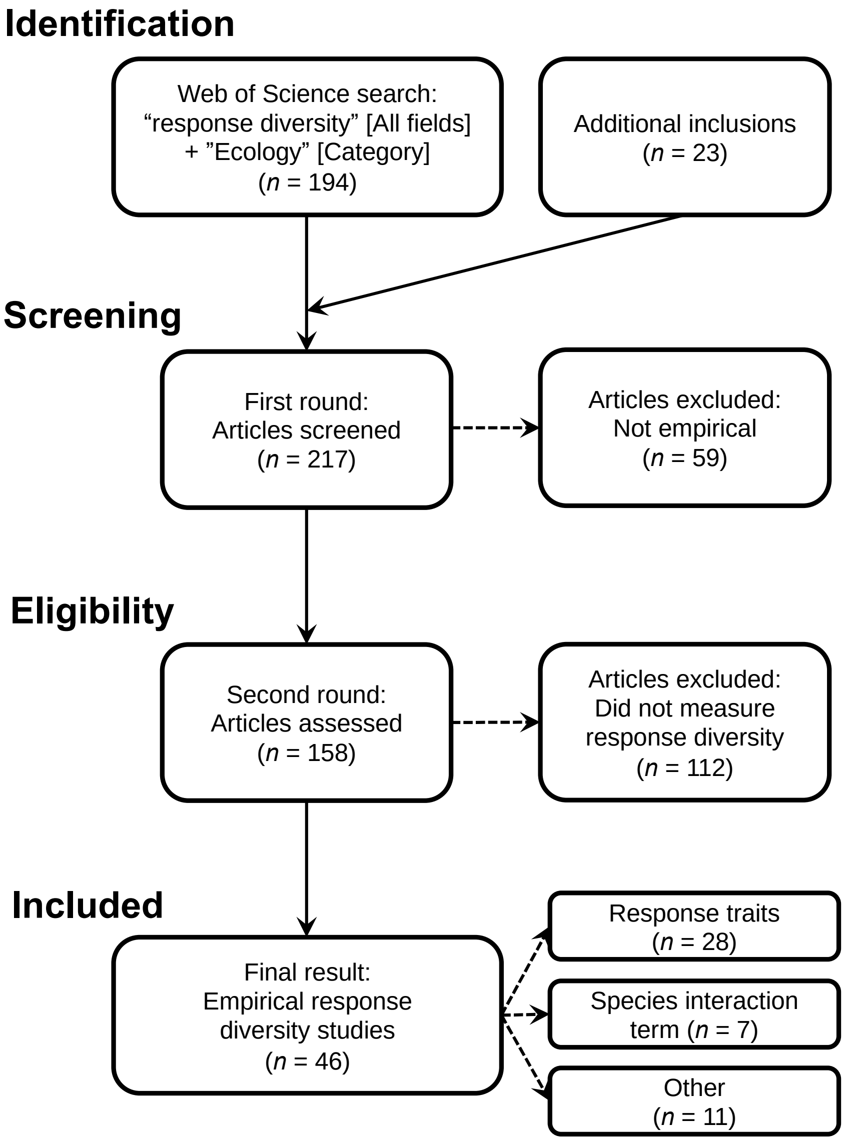


Supporting Figure S1. PRISMA diagram detailing the results of our systematic review. On 17 Dec 2021 (updated on 16 Jan 2023), we conducted a Web of Science search for "response diversity" in all fields within the "Ecology" Web of Science category. We supplemented the resulting 194 papers with 13 that were returned using the term "response variation" or that we otherwise knew to be relevant but were not included in the initial search results (*n* = 4), and on 23 Jan 2023, with 10 additional in press articles (not yet indexed in Web of Science) from journals which appeared more than 3 times in search results *(Glob. Chang. Biol.; J. Appl. Ecol.; Ecol. Soc.; Ecol. Appl.; Ecol. Lett.; Ecology; Ecol. Evol.; Ecography; Front. Ecol. Evol.; Ecosystems; Ecosphere; Proc. Roy. Soc. B; Mar. Ecol. prog. Ser.; J. Ecol.; Funct. Ecol.; Diversity; Divers. Dist.; Agri. Ecosyst. Environ.; Restor. Ecol.; J. Anim. Ecol.; Glob. Ecol. Biogeog.; Biol. Conserv.; Basic Appl. Ecol.*). We first excluded 59 articles that were not empirical studies (reviews, perspectives, *etc*.). Next, we screened the full-text articles and excluded 112 papers that did not measure response diversity (whether termed response diversity or not) based on the broad definition that response diversity should in some way characterise the range of species responses to environmental conditions. The resulting 46 empirical response diversity papers measured response diversity using various methods, the majority using functional response traits, followed by species-specific interaction terms in multi-species models of responses to the environment (see main text for further description of methods). For consistency in categorisation and exclusion criteria, all screening was conducted by a single researcher (S.R.P-J.R.).

**List of papers included in results after screening for eligibility.** Reference list is broken into screening categories from Figure S1.

*Response traits* (*n* = 28)

Aguirre-Gutiérrez, J., WallisDeVries, M. F., Marshall, L., van't Zelfde, M., Villalobos-Arámbula, A. R., Boekelo, B., Bartholomeus, H., Franzén, M., & Biesmeijer, J. C. (2017). Butterflies show different functional and species diversity in relationship to vegetation structure and land use. *Global Ecology and Biogeography*, *26*(10), 1126–1137. https://doi.org/10.1111/geb.12622

Altomare, M., Vasconcelos, H. L., Raymundo, D., Lopes, S., Vale, V., & Prado-Junior, J. (2021). Assessing the fire resilience of the savanna tree component through a functional approach. *Acta Oecologica*, *111*, 103728. https://doi.org/10.1016/j.actao.2021.103728

Aquilué, N., Filotas, É., Craven, D., Fortin, M.-J., Brotons, L., & Messier, C. (2020). Evaluating forest resilience to global threats using functional response traits and network properties. *Ecological Applications*, *30*(5), e02095. https://doi.org/10.1002/eap.2095

Bruno, D., Gutiérrez-Cánovas, C., Sánchez-Fernández, D., Velasco, J., & Nilsson, C. (2016). Impacts of environmental filters on functional redundancy in riparian vegetation. *Journal of Applied Ecology*, *53*(3), 846–855. https://doi.org/10.1111/1365-2664.12619

Carturan, B. S., Parrott, L., & Pither, J. (2022). Functional Richness and Resilience in Coral Reef Communities. *Frontiers in Ecology and Evolution*, *10*. https://www.frontiersin.org/articles/10.3389/fevo.2022.780406

Chillo, V., Anand, M., & Ojeda, R. A. (2011). Assessing the Use of Functional Diversity as a Measure of Ecological Resilience in Arid Rangelands. *Ecosystems*, *14*(7), 1168–1177. https://doi.org/10.1007/s10021-011-9475-1

Chillo, V., & Ojeda, R. (2014). Disentangling ecosystem responses to livestock grazing in drylands. *Agriculture, Ecosystems & Environment*, *197*, 271–277. https://doi.org/10.1016/j.agee.2014.08.011

Concepción, E. D., Moretti, M., Altermatt, F., Nobis, M. P., & Obrist, M. K. (2015). Impacts of urbanisation on biodiversity: The role of species mobility, degree of specialisation and spatial scale. *Oikos*, *124*(12), 1571–1582. https://doi.org/10.1111/oik.02166

Cooke, R. S. C., Bates, A. E., & Eigenbrod, F. (2019). Global trade-offs of functional redundancy and functional dispersion for birds and mammals. *Global Ecology and Biogeography*, *28*(4), 484–495. https://doi.org/10.1111/geb.12869

Correia, D. L. P., Raulier, F., Bouchard, M., & Filotas, É. (2018). Response diversity, functional redundancy, and post-logging productivity in northern temperate and boreal forests. *Ecological Applications*, *28*(5), 1282–1291. https://doi.org/10.1002/eap.1727

Craven, D., Filotas, E., Angers, V. A., & Messier, C. (2016). Evaluating resilience of tree communities in fragmented landscapes: Linking functional response diversity with landscape connectivity. *Diversity and Distributions*, *22*(5), 505–518. https://doi.org/10.1111/ddi.12423

Curzon, M. T., D'Amato, A. W., & Palik, B. J. (2016). Bioenergy harvest impacts to biodiversity and resilience vary across aspen-dominated forest ecosystems in the Lake States region, USA. *Applied Vegetation Science*, *19*(4), 667–678. https://doi.org/10.1111/avsc.12256

Döbert, T. F., Webber, B. L., Sugau, J. B., Dickinson, K. J. M., & Didham, R. K. (2017). Logging increases the functional and phylogenetic dispersion of understorey plant communities in tropical lowland rain forest. *Journal of Ecology*, *105*(5), 1235–1245. https://doi.org/10.1111/1365-2745.12794

Ford, H., Healey, J. R., Markesteijn, L., & Smith, A. R. (2018). How does grazing management influence the functional diversity of oak woodland ecosystems? A plant trait approach. *Agriculture, Ecosystems & Environment*, *258*, 154–161. https://doi.org/10.1016/j.agee.2018.02.025

Hordley, L. A., Gillings, S., Petchey, O. L., Tobias, J. A., & Oliver, T. H. (2021). Diversity of response and effect traits provides complementary information about avian community dynamics linked to ecological function. *Functional Ecology*, *35*(9), 1938–1950. https://doi.org/10.1111/1365-2435.13865

Laliberté, E., Wells, J. A., Declerck, F., Metcalfe, D. J., Catterall, C. P., Queiroz, C., Aubin, I., Bonser, S. P., Ding, Y., Fraterrigo, J. M., McNamara, S., Morgan, J. W., Merlos, D. S., Vesk, P. A., & Mayfield, M. M. (2010). Land-use intensification reduces functional redundancy and response diversity in plant communities. *Ecology Letters*, *13*(1), 76–86. https://doi.org/10.1111/j.1461-0248.2009.01403.x

Mandle, L., & Ticktin, T. (2015). Moderate land use changes plant functional composition without loss of functional diversity in India's Western Ghats. *Ecological Applications*, *25*(6), 1711–1724. https://doi.org/10.1890/15-0068.1

Mina, M., Messier, C., Duveneck, M., Fortin, M.-J., & Aquilué, N. (2021). Network analysis can guide resilience-based management in forest landscapes under global change. *Ecological Applications*, *31*(1), e2221. https://doi.org/10.1002/eap.2221

Morel, L., Barbe, L., Jung, V., Clément, B., Schnitzler, A., & Ysnel, F. (2020). Passive rewilding may (also) restore phylogenetically rich and functionally resilient forest plant communities. *Ecological Applications*, *30*(1), e02007. https://doi.org/10.1002/eap.2007

Mumme, S., Jochum, M., Brose, U., Haneda, N. F., & Barnes, A. D. (2015). Functional diversity and stability of litter-invertebrate communities following land-use change in Sumatra, Indonesia. *Biological Conservation*, *191*, 750–758. https://doi.org/10.1016/j.biocon.2015.08.033

Rader, R., Bartomeus, I., Tylianakis, J. M., & Laliberté, E. (2014). The winners and losers of land use intensification: Pollinator community disassembly is non-random and alters functional diversity. *Diversity and Distributions*, *20*(8), 908–917. https://doi.org/10.1111/ddi.12221

Rader, R., Birkhofer, K., Schmucki, R., Smith, H. G., Stjernman, M., & Lindborg, R. (2014). Organic farming and heterogeneous landscapes positively affect different measures of plant diversity. *Journal of Applied Ecology*, *51*(6), 1544–1553. https://doi.org/10.1111/1365-2664.12344

Sasaki, T., Lu, X., Hirota, M., & Bai, Y. (2019). Species asynchrony and response diversity determine multifunctional stability of natural grasslands. *Journal of Ecology*, *107*(4), 1862–1875. https://doi.org/10.1111/1365-2745.13151

Schnabel, F., Liu, X., Kunz, M., Barry, K. E., Bongers, F. J., Bruelheide, H., Fichtner, A., Härdtle, W., Li, S., Pfaff, C.-T., Schmid, B., Schwarz, J. A., Tang, Z., Yang, B., Bauhus, J., von Oheimb, G., Ma, K., & Wirth, C. (2021). Species richness stabilizes productivity via asynchrony and drought-tolerance diversity in a large-scale tree biodiversity experiment. *Science Advances*, *7*(51), eabk1643. https://doi.org/10.1126/sciadv.abk1643

Soria, M., Gutiérrez-Cánovas, C., Bonada, N., Acosta, R., Rodríguez-Lozano, P., Fortuño, P., Burgazzi, G., Vinyoles, D., Gallart, F., Latron, J., Llorens, P., Prat, N., & Cid, N. (2020). Natural disturbances can produce misleading bioassessment results: Identifying metrics to detect anthropogenic impacts in intermittent rivers. *Journal of Applied Ecology*, *57*(2), 283–295. https://doi.org/10.1111/1365-2664.13538

Spasojevic, M. J., Bahlai, C. A., Bradley, B. A., Butterfield, B. J., Tuanmu, M.-N., Sistla, S., Wiederholt, R., & Suding, K. N. (2016). Scaling up the diversity–resilience relationship with trait databases and remote sensing data: The recovery of productivity after wildfire. *Global Change Biology*, *22*(4), 1421–1432. https://doi.org/10.1111/gcb.13174

Thornhill, I., Biggs, J., Hill, M., Briers, R., Gledhill, D., Wood, P., Gee, J., & Hasall, C. (2018). The functional response and resilience in small waterbodies along land-use and environmental gradients. *Global Change Biology*, *in press*(March), 1–14. https://doi.org/10.1111/gcb.14149

Uhl, B., Wölfling, M., & Fiedler, K. (2021). Qualitative and Quantitative Loss of Habitat at Different Spatial Scales Affects Functional Moth Diversity. *Frontiers in Ecology and Evolution*, *9*. https://www.frontiersin.org/articles/10.3389/fevo.2021.637371

*Species interaction term* (*n* = 7)

Bartomeus, I., Park, M. G., Gibbs, J., Danforth, B. N., Lakso, A. N., & Winfree, R. (2013). Biodiversity ensures plant–pollinator phenological synchrony against climate change. *Ecology Letters*, *16*(11), 1331–1338. https://doi.org/10.1111/ele.12170

Cariveau, D. P., Williams, N. M., Benjamin, F. E., & Winfree, R. (2013). Response diversity to land use occurs but does not consistently stabilise ecosystem services provided by native pollinators. *Ecology Letters*, *16*(7), 903–911. https://doi.org/10.1111/ele.12126

Fründ, J., Zieger, S. L., & Tscharntke, T. (2013). Response diversity of wild bees to overwintering temperatures. *Oecologia*, *173*(4), 1639–1648. https://doi.org/10.1007/s00442-013-2729-1

Malyshev, A. V., Arfin Khan, M. A. S., Beierkuhnlein, C., Steinbauer, M. J., Henry, H. A. L., Jentsch, A., Dengler, J., Willner, E., & Kreyling, J. (2016). Plant responses to climatic extremes: Within-species variation equals among-species variation. *Global Change Biology*, *22*(1), 449–464. https://doi.org/10.1111/gcb.13114

McWilliam, M., Pratchett, M. S., Hoogenboom, M. O., & Hughes, T. P. (2020). Deficits in functional trait diversity following recovery on coral reefs. *Proceedings of the Royal Society B: Biological Sciences*, *287*(1918), 20192628. https://doi.org/10.1098/rspb.2019.2628

Stavert, J. R., Pattemore, D. E., Gaskett, A. C., Beggs, J. R., & Bartomeus, I. (2017). Exotic species enhance response diversity to land-use change but modify functional composition. *Proceedings of the Royal*

*Society B: Biological Sciences*, *284*(1860), 20170788–20170788. https://doi.org/10.1098/rspb.2017.0788

Winfree, R., & Kremen, C. (2009). Are ecosystem services stabilized by differences among species? A test using crop pollination. *Proceedings of the Royal Society B: Biological Sciences*, *276*(1655), 229–237. https://doi.org/10.1098/rspb.2008.0709

*Other* (*n* = 11)

Dell, J. E., Salcido, D. M., Lumpkin, W., Richards, L. A., Pokswinski, S. M., Loudermilk, E. L., O'Brien, J. J., & Dyer, L. A. (2019). Interaction Diversity Maintains Resiliency in a Frequently Disturbed Ecosystem. *Frontiers in Ecology and Evolution*, *7*, 145. https://doi.org/10.3389/fevo.2019.00145

Dzubakova, K., Peter, H., Bertuzzo, E., Juez, C., Franca, M. J., Rinaldo, A., & Battin, T. J. (2018). Environmental heterogeneity promotes spatial resilience of phototrophic biofilms in streambeds. *Biology Letters*, *14*(10), 20180432. https://doi.org/10.1098/rsbl.2018.0432

Fauchald, P., Skov, H., Skern-Mauritzen, M., Hausner, V. H., Johns, D., & Tveraa, T. (2011). Scale-dependent response diversity of seabirds to prey in the North Sea. *Ecology*, *92*(1), 228–239. https://doi.org/10.1890/10-0818.1

Feit, B., Blüthgen, N., Daouti, E., Straub, C., Traugott, M., & Jonsson, M. (2021). Landscape complexity promotes resilience of biological pest control to climate change. *Proceedings of the Royal Society B: Biological Sciences*, *288*(1951), 20210547. https://doi.org/10.1098/rspb.2021.0547

Joseph, G. S., Seymour, C. L., Cumming, G. S., Cumming, D. H. M., & Mahlangu, Z. (2014). Termite Mounds Increase Functional Diversity of Woody Plants in African Savannas. *Ecosystems*, *17*(5), 808–819. https://doi.org/10.1007/s10021-014-9761-9

Leary, D. J., & Petchey, O. L. (2009). Testing a biological mechanism of the insurance hypothesis in experimental aquatic communities. *Journal of Animal Ecology*, *78*(6), 1143–1151. https://doi.org/10.1111/j.1365-2656.2009.01586.x

McCann, M. J. (2016). Response diversity of free-floating plants to nutrient stoichiometry and temperature: Growth and resting body formation. *PeerJ*, *4*, e1781. https://doi.org/10.7717/peerj.1781

Moore, J. W., & Olden, J. D. (2017). Response diversity, nonnative species, and disassembly rules buffer freshwater ecosystem processes from anthropogenic change. *Global Change Biology*, *23*(5), 1871–1880. https://doi.org/10.1111/gcb.13536

Sanford, M. P., Manley, P. N., & Murphy, D. D. (2009). Effects of urban development on ant communities: Implications for ecosystem services and management. *Conservation Biology*, *23*(1), 131–141. https://doi.org/10.1111/j.1523-1739.2008.01040.x

Schweiger, A. H., Audorff, V., & Beierkuhnlein, C. (2015). The acid taste of climate change: 20th century acidification is re-emerging during a climatic extreme event. *Ecosphere*, *6*(6), art94. https://doi.org/10.1890/ES15-00032.1

Thibaut, L. M., Connolly, S. R., & Sweatman, H. P. A. (2012). Diversity and stability of herbivorous fishes on coral reefs. *Ecology*, *93*(4), 891–901. https://doi.org/10.1890/11-1753.1
